# Postnatal conversion of methylcytosine to hydroxymethylcytosine reconfigures the human neuronal epigenome

**DOI:** 10.64898/2026.02.18.706675

**Authors:** Heng Xu, Jo-Fan Chien, Alexey Kozlenkov, Ramu Vadukapuram, Junhao Li, Yu Wei, Andrew J. Dwork, Chunyu Liu, Stella Dracheva, Eran A. Mukamel

## Abstract

Healthy brain development requires a coordinated process of postnatal cellular maturation throughout the first two decades of life that transforms neuronal morphology, connectivity, physiology, and gene expression. The maturation and stable maintenance of neuron identity is driven, in part, by large-scale reconfiguration of the neuronal DNA methylome. Neurons have uniquely high levels of 5-hydroxy-methyl-cytosine (hmC) compared to other cell types, yet the relative contributions of 5hmC and 5-methyl-cytosine (mC) remain unknown because most experimental assays do not distinguish these marks. We measured mC and hmC using bisulfite- and oxidative-bisulfite sequencing in excitatory and inhibitory neurons, along with mRNA and histone modifications, from the prefrontal cortex of 103 human donors, ranging from 38 days to 77 years of age. Up to half of all CG dinucleotides convert from mC to hmC in a gradual process extending throughout the first decade of life, dramatically reshaping the neuronal methylome. Asymmetric enrichment of hmC on the sense strand of actively transcribed genes increases in a linear, clock-like fashion throughout the lifespan, indicating a mechanistic link between transcription and hmC. We found that sex differences in X-linked DNA methylation in the human brain are primarily driven by hmCG rather than mCG, suggesting an important role for hmC in X-chromosome inactivation (XCI) and escape gene expression. We found key changes in 5hmC at dynamic cis-regulatory elements marked by changing cell type-specific levels of active and repressive histone modifications. Collectively, our findings reveal the dynamic trajectory of hmC in human neurons across the lifespan and highlight the association of DNA hydroxymethylation with transcription, chromatin state, and sex-specific gene regulation.

## Introduction

The landscape of DNA methylation in neurons changes dramatically throughout the lifespan as cells establish and maintain their unique epigenetic identity^1–4^. The configuration of DNA methylation in neurons differs from most other cell types in two ways, both of which emerge gradually over the first two decades of post-natal development in humans. Methylation at CA and other non-CG dinucleotides, collectively called mCH, accumulates to become the dominant contributor of methylcytosine in adult neurons, encoding a cell type-specific signature of gene regulation^5–9^. At the same time, a parallel process converts a subset of methylcytosines, mainly at CG sites, to hydroxymethylcytosine (hmC) via TET enzymes^10–15^. Both hmC and mCH have been linked with potential roles in neuronal function^12,16–20^, implicated in neurodevelopmental disorders including Rett’s syndrome and Beck-Fahrner syndrome^21,22^, and associated with neurodegenerative Alzheimer’s disease^23,24^. Yet, the lifelong dynamics of hmC in specific human neuron types and their relation with other epigenetic marks and gene expression remain largely unknown.

One model proposes that hmC acts mainly as an intermediate in the process of active demethylation. According to this hypothesis, hmC is part of a closed cycle in which C converts to mC, then to hmC, and returns to unmethylated C via one of several potential pathways^25,26^. This model predicts hmC is a transient modification, whose presence in the adult brain indicates a dynamical steady-state. Alternatively, the abundant hmC could constitute a stable form of functional demethylation that disrupts the repressive effect of methylcytosine on transcription factor binding and gene expression, independent of active demethylation^12^. In either scenario, the dynamic interconversion of C, mC and hmC from infancy through adolescence and adulthood could be critical for healthy cognitive development and neural function^14,27^.

Advances in single cell bisulfite sequencing and multi-ome technology have revealed a highly cell type-specific landscape of total DNA methylation (i.e. mC+hmC)^28–31^, but measurements of hmC with single cell resolution have been relatively limited^32–35^. The cell-type specific dynamics of hmC in the developing human brain remain unexplored. To address these gaps, we performed whole-genome bisulfite and oxidative bisulfite sequencing (BS-seq, OxBS-seq) to map mC and hmC in glutamatergic (Glu) and medial ganglionic-eminence derived GABAergic (MGE-GABA) neurons from the human prefrontal cortex^36,37^. We contextualize these data with matched gene expression and histone modification measurements in the same cell types and ages. Our multimodal dataset includes post-mortem samples from 103 male and female donors ranging in age from infancy to >70 years old, covering key milestones and transitions in brain development. We describe and integrate distinct age-related changes in DNA methylation, histone modifications, and gene expression, offering insights into the regulatory mechanisms that shape neuronal function and plasticity across the human lifespan. Our data are available for browsing and analysis as a resource at https://brainome.ucsd.edu/BrainHydroxyDevelopment.

## Results

### Genome-wide conversion of mC to hmC during human postnatal brain development

To assess the dynamic trajectory of mC and hmC throughout postnatal brain development, we collected samples of human prefrontal cortex (Brodmann area BA9) from 103 donors ranging from 38 days to 77 years old (Fig. 1A,B). Neuron types have highly specific epigenetic and transcriptomic signatures that are evident in single cell sequencing data^8,38^. However, existing single-cell and -nucleus methods for measuring hmCG remain challenging, particularly for profiling large numbers of neurons^34,39^. We therefore applied multicolor fluorescence-activated nuclei sorting (FANS) to isolate two major neuronal populations: excitatory glutamatergic neurons (Glu, NeuN+/SOX6-) and medial ganglionic eminence-derived inhibitory GABAergic interneurons (MGE-GABA, NeuN+/SOX6+)^37^. The MGE-GABA neurons include the parvalbumin (PV) and somatostatin (SST) expressing populations^36,37,40^, while the Glu population consists mainly of excitatory neurons but also includes a small proportion of non-MGE-derived GABAergic neurons that do not express SOX6, such as VIP and LAMP5 expressing cells. We used RNA sequencing (RNA-seq) to assess gene expression, whole genome bisulfite sequencing (BS-seq) to quantify total modified cytosine (total mC = mC + hmC), and oxidative bisulfite sequencing (OxBS-seq) to measure methylcytosine (mC)^41^. Hydroxymethylcytosine (hmC) was estimated using the difference (total mC - mC) at each site. We further used chromatin immunoprecipitation sequencing (ChIP-seq) to profile post-translational histone modifications at active enhancers and promoters (H3K27ac), polycomb repressive complex 2 (PRC2)-mediated repressive regions (H3K27me3), and constitutive heterochromatin (H3K9me3). Our dataset is an unprecedented and comprehensive survey of human postnatal brain development comprising 116 transcriptomes, 338 methylomes (12-15x genome coverage per sample), and 131 ChIP-seq datasets, with more than 80 billion total sequenced reads (Table S1, Fig. S1A).

**Figure 1.**
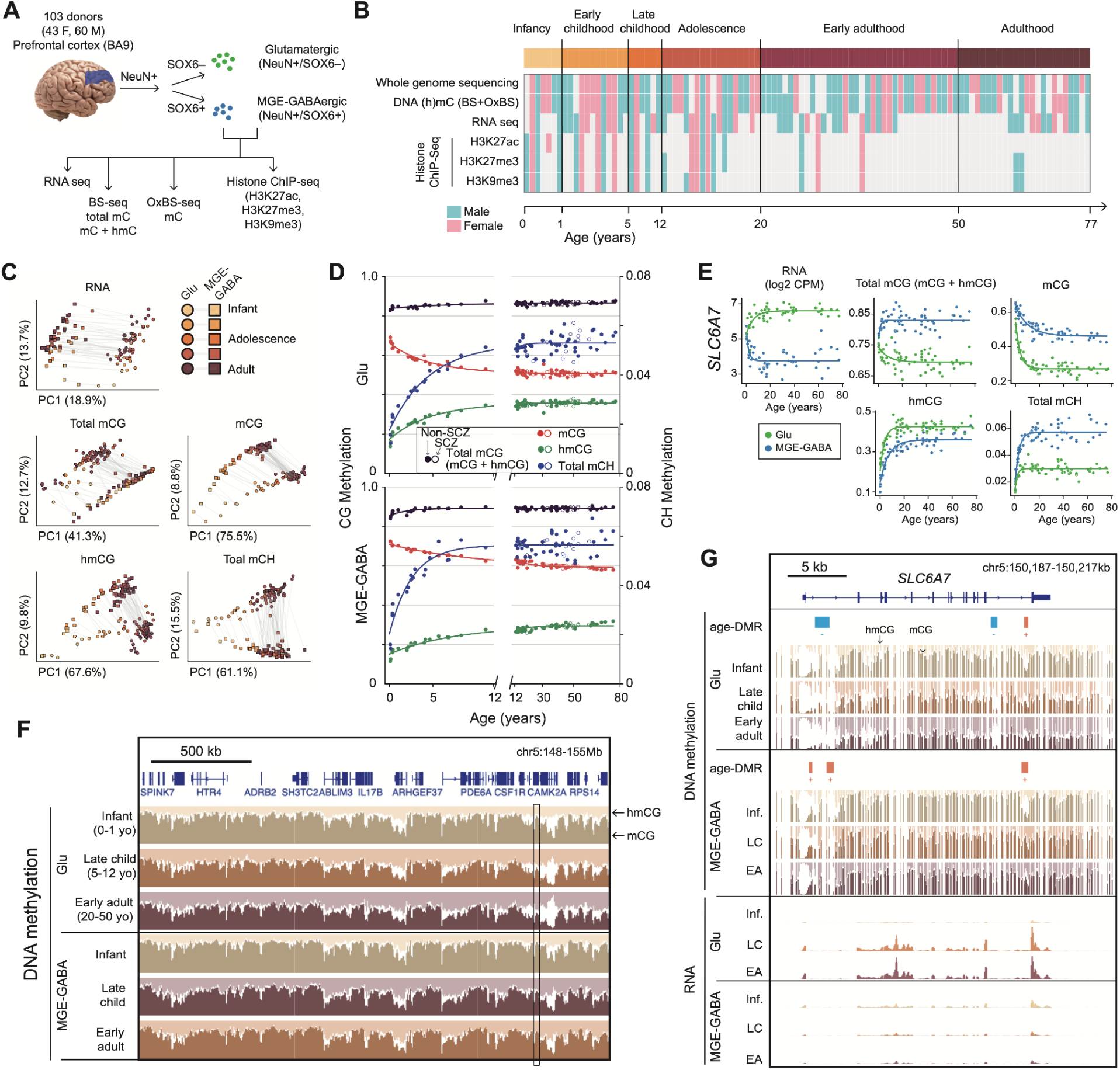
Mapping dynamic changes of genome-wide DNA (hydroxy)methylation and gene expression in two major neuron types across the lifespan. (A,B) Cell type-specific transcriptomic and epigenomic sequencing from Brodmann area BA9 from 103 postmortem donors aged 0-77 years. (C) Principal component (PC) analysis of RNA expression, gene body total mCG, mCG, hmCG and total mCH levels across age groups and neuronal classes. Gray lines connect samples from the same donor. During postnatal development, Glu and MGE-GABA neurons have increasingly distinct mCG, hmCG and mCH patterns; RNA and total mCG patterns are less distinct. (D) Genome-wide average methylation levels across age. SCZ: schizophrenia. (E) RNA expression and gene body methylation of the proline transporter, *SLC6A7,* in Glu and MGE-GABA neurons. (F,G) Browser view of DNA methylome tracks and transcriptome (RNA-seq) around *SLC6A7* locus in Glu and MGE-GABA neurons. For DNA methylation, the mC level at each CG position is plotted in a darker color extending up from the bottom of the track, while hmC is shown in a lighter color extending down from the top of the track. The proportion of unmethylated cytosine corresponds to the gap between the mC and hmC levels. Boxed region in (F) corresponds to the locus shown in (G). Age-DMRs: Differentially methylated regions (+/-: increasing/decreasing mCG with age).

Across subjects, DNA methylomes and transcriptomes were highly consistent (Fig. S1B). The correlation of gene expression and of gene body methylation levels between subjects of the same age were high (Spearman r>0.95), and were slightly lower across age groups (>0.93). The correlations between neuronal classes (Glu versus MGE-GABA) within each age group were notably lower (RNA: 0.88 ± 0.02; total mCG: 0.93 ± 0.01; mCG: 0.92 ± 0.01; mCH: 0.87 ± 0.04), consistent with the cell type specificity of transcriptomic and epigenomic signatures. Glu and MGE-GABA neurons from infants were less correlated with other age groups, and more correlated with each other, indicating a similar immature neuronal state for both lineages (Fig. S1C).

To assess the post-natal developmental trajectory of neurons, we performed principal component (PC) analysis on gene body DNA methylation and on gene expression (Fig. 1C). Notably, the diverging trajectories of mCG, hmCG and mCH for Glu vs. MGE-GABA cells reflect increasing differentiation of the major neuronal populations with age. By contrast, RNA and total mCG trajectories did not clearly follow this pattern, highlighting the need to distinguish mC and hmC dynamics. Gene expression was most dynamic early in postnatal development, while cell type-specific DNA methylation emerged more gradually.

Previous studies showed that hmCG is largely absent in fetal and early infant brain^5,13,16^, yet the dynamic trajectory and functional significance of the global accumulation of hmCG in specific neuron types remain unknown. As in our earlier study in adult donors^37^, we found that genome-wide hmCG was most enriched in adult Glu neurons (∼37% of all CG dinucleotides) compared with MGE-GABA (30%), while mCG had the opposite pattern (44% in Glu, 54% in MGE-GABA, Fig. S1D)^37^. We found that this high level of hmCG in neurons arises gradually over the first two decades of life in an extended process of conversion of mCG to hmCG (Fig. 1D). In contrast, the total mCG level (mCG + hmCG) changed by less than 3% in both cell types, suggesting that mCG loss is largely balanced by hmCG gain on average throughout the genome. By fitting a simple first-order dynamical model (Methods), we found that the genome-wide loss of mCG proceeds faster in Glu neurons (exponential time constant τ = 4.5 ± 0.4 years, SE) than in MGE-GABA neurons (10.1 ± 0.8 yrs). Likewise, hmCG accumulates faster in Glu neurons (4.8 ± 0.4 yrs) compared with MGE-GABA neurons (7.3 ± 0.6 yrs). In contrast, mCH increases more rapidly in MGE-GABA neurons (2.6 ± 0.3 yrs) compared with Glu neurons (4.7 ± 0.4 yrs).

The levels of mCG and hmCG are tightly linked with the functional roles of specific genomic compartments (Fig. S2A). Active regions, including gene bodies (exons and introns), promoters without CG islands (CGI), and enhancers, had more hmCG compared to repressed regions, including H3K27me3-marked domains and LINE-1 retrotransposons (L1HS) (Fig. S2A). CGI promoters and DNA methylation valleys had low levels of both mCG and hmCG. Notably, Glu neurons accumulated more hmCG in repressed regions than MGE-GABA neurons (Fig. S2A). Conversely, MGE-GABA enhancers accumulated more hmCG than Glu enhancers.

The cell type–specific dynamic trajectories of mC and hmC could potentially reflect differences in the activity of DNA methyltransferase and/or TET enzymes. However, we found no direct correlation between DNA methylation dynamics and the age-dependent expression patterns of the three TET genes, which catalyze the conversion of mC to hmC^10,11^, or the DNA methyltransferases responsible for de novo (DNMT3A) and maintenance methylation (DNMT1)^42^ (Fig. S2B).

In addition to hmCG, neurons accumulate substantial non-CG methylation (mCH), an epigenetic mark that is not detected at high levels in other body tissues^5,7,43^. We observed a progressive increase in mCH during postnatal development, rising from 2% to 5% of CH sites in Glu neurons and to 6% in MGE-GABA neurons (Fig. S1D). This increase was largely driven by mCA, which increased from about 2% to ∼10% (Fig. S1E). Hydroxymethylation at both CG and CA dinucleotides can enable binding of MECP2^12,21^, raising the question of whether hmCH also plays a major role in neuronal regulatory dynamics. We found that non-CG hydroxymethylation (hmCH) remains below 1.1% in both neuron types (Fig. S1D)^17,37^.

Our data provide fine-grained, base-resolution information about cell type-specific gene regulation across human postnatal brain development. For example, *SLC6A7*, encoding the sodium-dependent high affinity proline transporter (PROT1) that is important for locomotion and learning^44,45^, has a similar expression level in Glu and MGE-GABA neurons during infancy. Over the first decade of life, *SLC6A7* expression is specifically upregulated in Glu and downregulated in MGE-GABA neurons (Fig. 1E). Consistent with this cell type difference in expression of *SLC6A7*, we found a more pronounced decrease in promoter and gene body mCG in Glu neurons compared to MGE-GABA neurons (Fig. 1E-G). We observed diverse trajectories of DNA methylation across genes, including those encoding key synapse-related proteins (e.g. *CAMK2A, SLC17A7)* and developmental factors (*DPYSL3*) (Fig. S3).

### Strand asymmetry of hmCG increases with age in expressed genes

During transcription, the DNA strands separate, allowing the antisense strand to serve as a template for mRNA synthesis. The different roles of the sense and antisense strands could lead to asymmetric impacts of gene expression on epigenetic marks. If so, this asymmetry should be most apparent in highly transcribed genes for which the sense strand (non-template) may be more accessible to DNA methyltransferase or TET enzymes. Consistent with this, we found that hmCG is enriched on the sense strand, and mCG on the antisense strand, within gene bodies of highly expressed genes in adult Glu and MGE-GABA neurons (Fig. 2A, S4A-B), as reported previously^36,46^. Non-expressed genes had no detectable asymmetry, and the level of asymmetry correlated with gene expression in a graded fashion. We found up to ∼3% asymmetry between the sense and antisense strands in mCG and hmCG throughout the gene body of adult samples. The asymmetry persisted downstream of the gene for several kb past the 3’ transcription end site. Highly expressed genes also had a slightly asymmetric (<0.2%) enrichment of total mCH on the antisense strand, which was absent in non-expressed genes (Fig. S4C).

**Figure 2.**
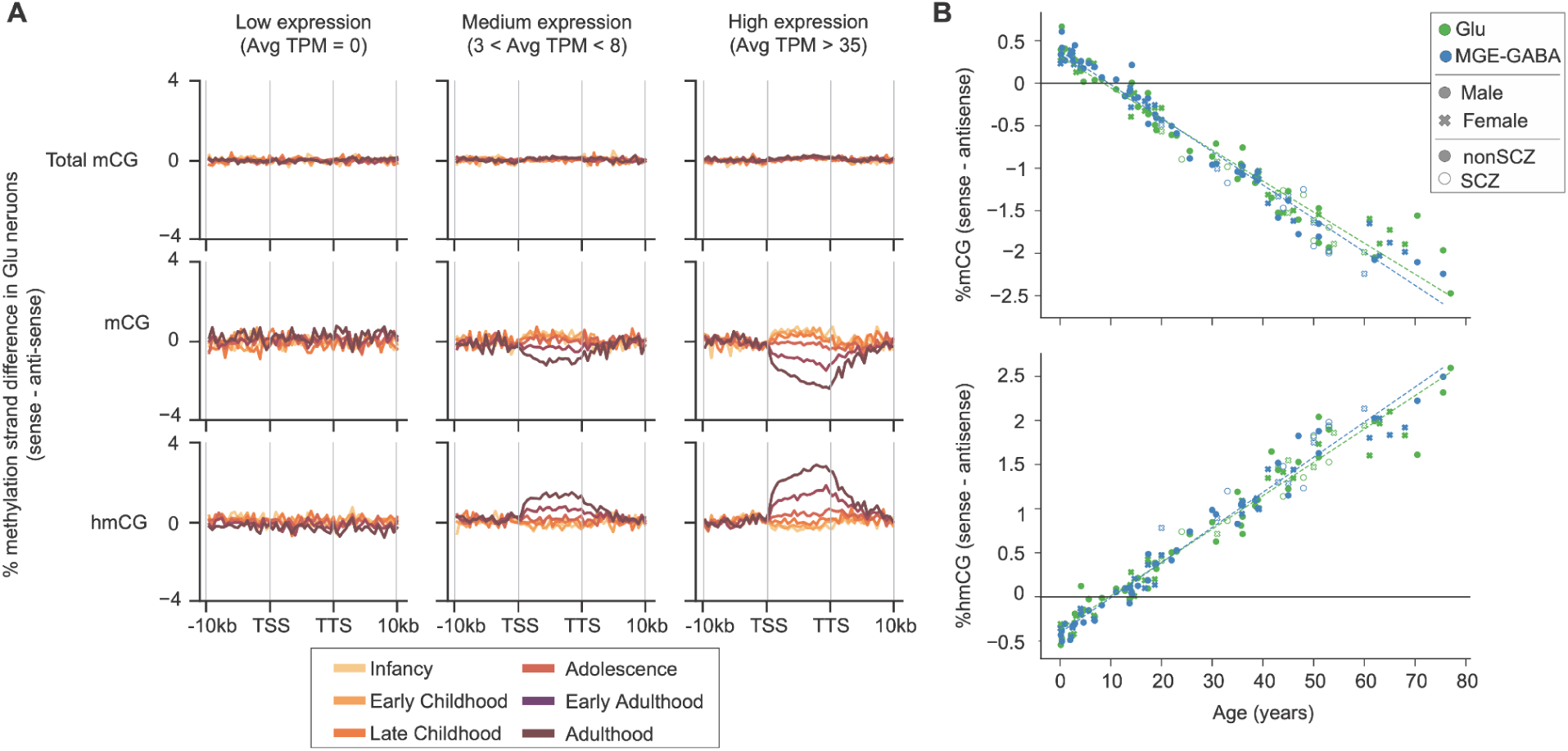
Clock-like accumulation of transcription-dependent strand asymmetry in gene body methylation. **(A)** Strand asymmetry (sense - antisense) of mCG and hmCG for low (average TPM = 0), medium (TPM 3-8) and highly expressed (TPM > 35) genes across age groups in Glu neurons. TSS/TTS, transcription start/termination site. **(B)** Average mCG and hmCG strand asymmetry within the gene body of highly expressed genes (top 10% across all age groups) in Glu and MGE-GABA. Linear regressions for each cell type are shown as dashed lines.

Strikingly, we found that the strand asymmetry in mCG and hmCG increases with age in a remarkably linear fashion across both neuronal classes (Pearson |r| = 0.97-0.98, p<1e-10; Fig. 2B, S4A). This correlation was equal to or stronger than that of DNA methylation-based clocks^47–51^ (e.g. r=0.935-0.991 for human brain^51^). The linear increase of hmCG strand asymmetry accurately predicts age (RMSE = 3.89-3.96 years), but unlike previous clocks it does not require fitting hundreds of adjustable regression weights for individual CG sites throughout the genome.

In infants, the strand asymmetry was small and had the opposite sign from adults, with higher mCG and lower hmCG on the sense compared with the antisense strand. The strand asymmetry increased by ∼0.3% per decade in both neuron types for the most highly expressed genes. These data suggest that transcription promotes conversion of mCG to hmCG on the sense strand. The asymmetric impact of mRNA expression, operating continuously throughout the lifespan, leaves a clock-like signature of chronological age encoded in the strand asymmetry of DNA methylation.

We found that thousands of individual genes had a significant correlation between mCG and/or hmCG strand asymmetry and age (Fig. S4D,E). The vast majority (83-93%) had a negative association of age with mCG asymmetry (sense - antisense) and a positive association with hmCG asymmetry. Notably, most genes with significant age-associated mCG and/or hmCG strand asymmetry were strongly expressed (TPM>12) (63-71%).

To determine whether dynamic, age-related gene expression alters the accumulation of asymmetric DNA methylation, we separately analyzed genes with significantly increasing (age-up) or decreasing (age-down) expression over the lifespan. There was no significant difference in the strand asymmetry between age up- versus down-regulated genes (Fig. S4F).

Notably, we observed no strand asymmetry in the level of total mCG (mCG+hmCG)^46^ (Fig. 2A). This suggests the strand asymmetry reflects TET-dependent conversion of mCG to hmCG, but not subsequent demethylation. Strand asymmetry is thus not detectable by conventional BS-seq alone, highlighting the importance of our OxBS and BS-seq data for measuring hmCG and mCG separately.

### Sex-specific neuronal DNA methylation on the X chromosome

DNA methylation plays a critical role in the process of X-chromosome inactivation (XCI) in females, ensuring equivalent expression of most X-linked genes in males and females (dosage compensation)^52^ (Fig. 3A). In mouse cerebellum, hmC is relatively depleted on chrX in both males and females^13^. However, the role of hmC in the dynamic regulation of XCI and dosage compensation has not been addressed in the human brain. We confirmed that autosomes from adult neurons had highly consistent levels of mC and hmC in males and females (Fig. 3B-F; Fig. S5A). Notably, chr19 had ∼2-fold lower total mCH compared to other autosomes, which may reflect the unique genomic features of human chr19 such as its high gene density and GC content^53^.

**Figure 3.**
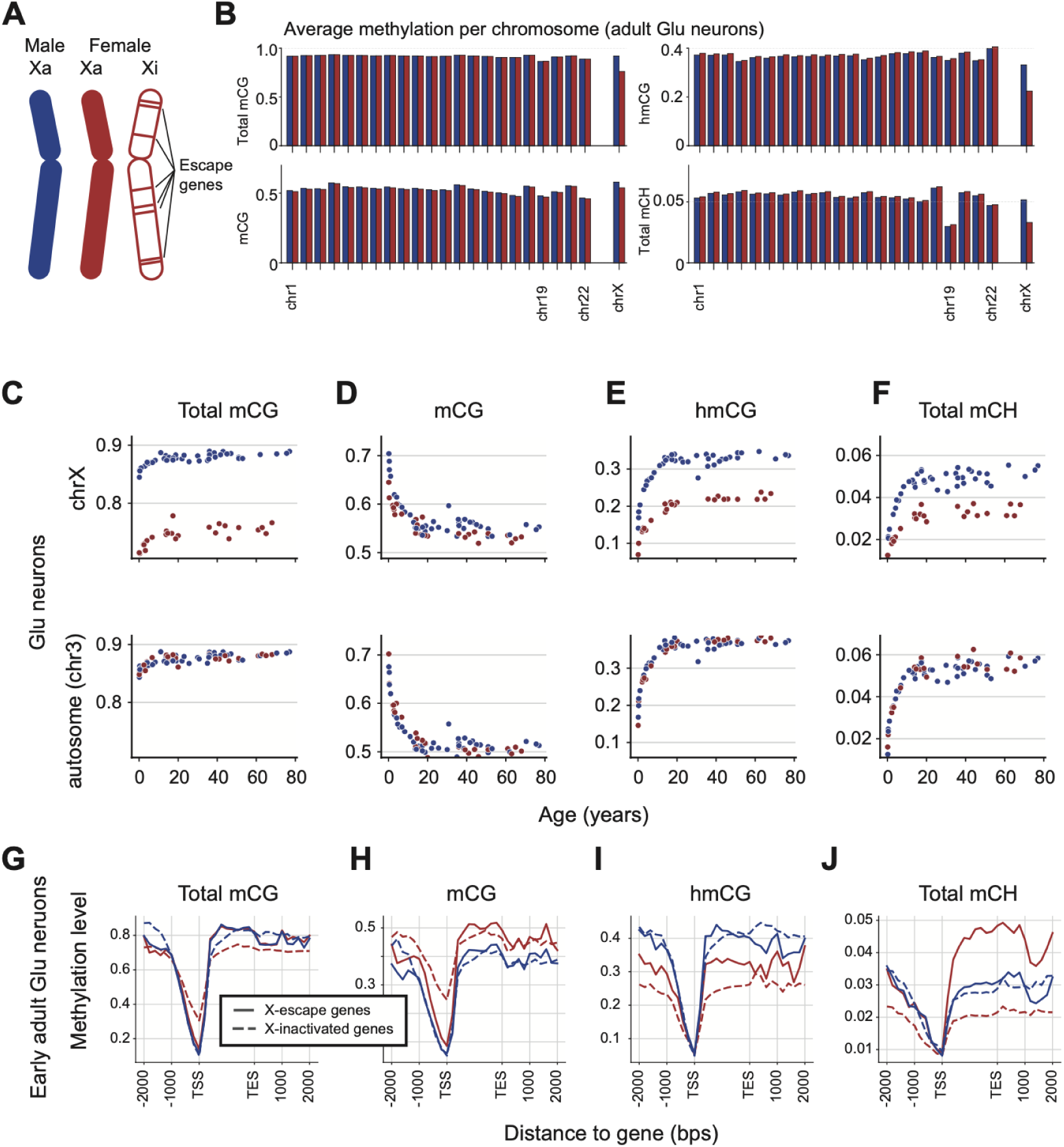
Contribution of hmC to female neuron-specific DNA methylation on chrX. **(A)** Illustration of active (Xa, solid color) and inactive (Xi, white) chrX alleles in males and females. **(B)** Average DNA methylation level per chromosome in adult Glu neurons, showing female-specific reduction in hmC and mCH on chrX. **(C–F)** Average DNA methylation level on chrX and on an autosome (chr3) in male and female Glu neurons across the lifespan. Sex differences in chrX hmCG and mCH arise early in brain development. **(G–J)** Methylation levels in X-inactivated and X-escape genes^54^ during early adulthood. The transcription start site (TSS) of inactivated genes has higher mCG in females, while gene bodies and flanking intergenic regions have reduced hmCG and mCH. Escape genes have increased hmCG and mCH.

In contrast with the autosomes, we found substantial sex differences on chrX. In both Glu and MGE-GABA neurons, females had ∼15% lower total mCG (mCG+hmCG) across chrX compared to males, consistent with reduced methylation on the inactive X chromosome (Xi) outside promoter regions^55,56^ (Fig. 3B,C, S5A). Notably, in Glu neurons, this difference was driven by 12% lower hmCG in females on average (Fig. 3E), whereas mCG levels were similar in males and females (Fig. 3D). In contrast, MGE-GABA neurons had lower mCG and hmCG in females (Fig. S5A). In both Glu and MGE-GABA neurons, mCH levels on chrX were approximately twice as high in adult males compared to females, consistent with the nearly complete absence of mCH on the inactive X chromosome (except at genes that escape X inactivation) in female mouse brain^55^ (Fig. 3E,F, S5A).

The lower level of hmCG in female chrX was already established in the youngest infants we examined (<2 months), prior to the postnatal onset of mCG to hmCG conversion (Fig. 3E). These data suggest that the inactive X chromosome in females may be selectively protected from TET-mediated oxidative conversion, similar to the protection of Xi from DNMT3A-dependent mCH accumulation in the mouse brain^55^.

A subset of ∼10-15% of X-linked genes in humans escape from XCI and are expressed from both X chromosomes in females^54,57^. Previous studies analyzing total mCG and mCH showed that these escape genes have distinct allele-specific DNA methylation patterns in neurons^5,7,55^. The promoters of most genes, which are subject to XCI, are hypermethylated on the Xi in females, whereas escape gene promoters remain hypomethylated on both alleles^55,58,59^. By distinguishing mCG and hmCG, we found that female-specific hypermethylation at promoters of genes subject to XCI is primarily driven by mCG, rather than hmCG (Fig. 3G-I). This is notably different from the pattern across the rest of the X chromosome, where sex differences in DNA methylation were predominantly attributable to hmCG (Fig. 3D,E). Furthermore, we observed less accumulation of hmCG near inactivated genes in females compared to males (Fig. 3I), suggesting that the TET-mediated conversion of mC to hmC may be less effective in densely packed chromatin regions such as Xi. Consistent with previous studies, we found that escape genes had higher gene body mCH in females^5,7,55^ (Fig. 3J). Thus, our data indicate that X chromosome DNA methylation differences between males and females, as well as between inactivated and escape genes, are established early in development, persist throughout the lifespan, and are consistent in both neuronal classes (Fig. 3C-F, Fig. S5B).

In contrast to the sexual dimorphism in X-linked DNA methylation, we expect that gene expression in male and female neurons is largely balanced throughout the lifespan except at a small number of genes specifically involved in XCI (sex-DEGs; Fig. S6A, Table S2). We identified 13 X-linked sex-DEGs (FDR < 0.05) in at least one neuronal class (11 in Glu; 13 in MGE-GABA), including the XCI master regulator non-coding RNA, *XIST*. Most of the X-linked sex-DEGs were classified as escapees in previous studies (11/13)^54^. Three male-biased genes, *ASMTL*, *GTPBP6,* and *PLCXD1*, are located in the pseudoautosomal region^60,61^. Additionally, the X-linked gene *GTPBP6* was more expressed in males, but its expression increases with age in both males and females (Fig. S6A).

### Dynamic methylation and demethylation across the lifespan

Using our high-coverage DNA methylomes, we identified age-related differentially methylated regions (age-DMRs) through pairwise comparisons among age groups from BS-seq data in each cell type. We report age-DMRs identified from BS-seq (which correspond to total mCG changes) rather than OxBS-seq (mCG) because, as shown above, the age-related conversion of mCG to hmCG occurs throughout the genome, and does not specifically mark discrete regulatory regions. Our analysis focuses on regions where mCG changes were not fully compensated by opposing hmCG changes, resulting in a net change in total mCG. These age-DMRs are therefore directly comparable with regions reported in previous BS-seq studies^29,31^.

We combined DMRs identified in all pairwise age group comparisons (Fig. S7A) and further filtered them to include regions with a substantial difference (>0.3) in mCG across age groups. We found 146,966 age-DMRs in Glu and 72,432 in MGE-GABA, covering 3.4% and 1.4% of the genome, respectively (Fig. S7B,C). Notably, the vast majority (87%) of our age-DMRs were identified as cell type-DMRs in single nucleus DNA methylomes of human neurons^30^, highlighting their likely role in cell type-specific epigenetic regulation. Age-DMRs also overlapped DMRs identified in previous studies of prenatal human brain development^31^ (31% of age-DMRs) and of aging^29^ (11%).

To quantify methylation dynamics within age-DMRs, we developed a three-state kinetic model guided by the known enzymatic pathways of DNA methylation and active demethylation (see Methods, Fig. 4A–C). Our model assumes that each genomic cytosine can exist in an unmethylated (C), methylated (mC) or hydroxymethylated (hmC) state. Three first-order kinetic pathways enable transitions from C to mC (mediated by DNMT3A), with time constant (inverse rate) τ_𝐶→𝑚𝐶_ ; from mC to hmC (mediated by TET), τ_𝑚𝐶→ℎ𝑚𝐶_ ; and from hmC to C (active demethylation), τ_ℎ𝑚𝐶→𝐶_ (Fig. 4C). For each DMR, we fit five model parameters (three time constants and two initial conditions specifying the abundance of mC and hmC at birth) using a least-squares criterion to provide a simple yet highly accurate description of the dynamic relationships among these marks throughout the lifespan. In terms of the time constants, the adult (steady state) methylation levels predicted by the model are: 𝑚𝐶_𝐴𝑑𝑢𝑙𝑡_ = τ_𝑚𝐶→ℎ𝑚𝐶_ /τ, ℎ𝑚𝐶_𝐴𝑑𝑢𝑙𝑡_ = τ_ℎ𝑚𝐶→𝐶_ /τ, where τ = τ_𝐶→𝑚𝐶_ + τ_𝑚𝐶→ℎ𝑚𝐶_ + τ_ℎ𝑚𝐶→𝐶_ . These dynamics can also be visualized using a ternary plot showing the initial and final (asymptotic) methylation levels as points in a triangular simplex, with a dynamic trajectory connecting the two points (Fig. 4B).

**Figure 4.**
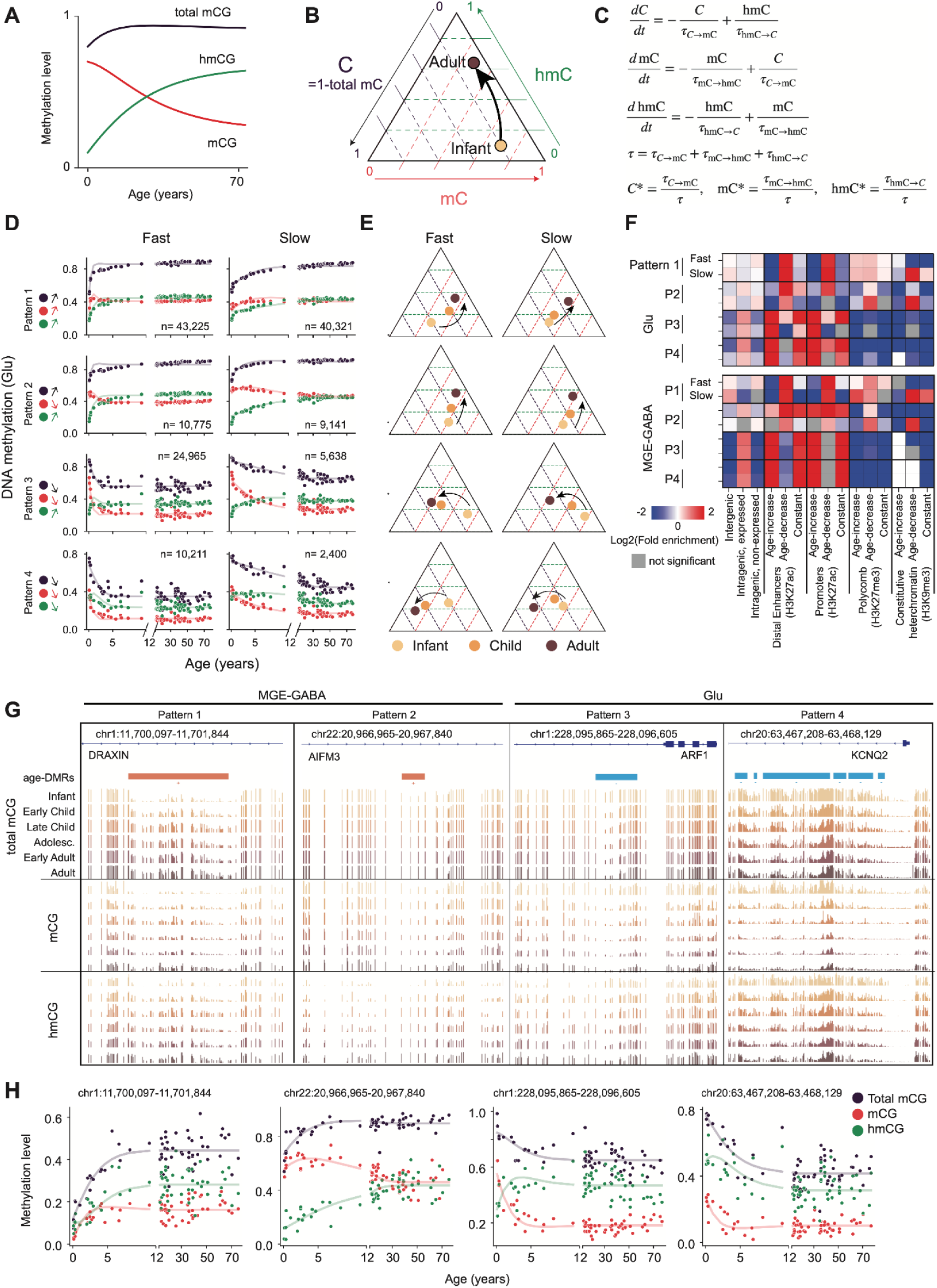
Dynamics of DNA methylation across the lifespan. **(A-C)** Schematic illustration of a 3-state model describing DNA methylation dynamics. (**A**) Example trajectories for total mCG, mCG, and hmCG. **(B)** Ternary plot showing the same trajectory as in (A), highlighting the interrelationship among the three forms of genomic cytosine. (**C**) Equations defining the 3-state kinetic model. C*/mC*/hmC* indicates the steady-state value for each variable, which corresponds to the fixed point attractor of the dynamical system (adult state). **(D)** Average DNA methylation profiles of age-DMRs grouped into eight patterns based on distinct methylation dynamics (see Methods) in Glu neurons. **(E)** Ternary plots of the four main age-DMR patterns (fast and slow) in Glu. **(F)** Enrichment of Glu age-DMRs in genomic regions (FDR < 0.05). Distal enhancers, promoters, Polycomb-repressed regions (H3K27me3), and constitutive heterochromatin (H3K9me3) are assigned based on our ChIP-seq data into age-activated, age-deactivated, and stable peaks. **(G)** Browser view of example age-DMRs, showing (left to right): patterns 1, 2 (shown in MGE-GABA), patterns 3 and 4 (shown in Glu). **(H)** DNA methylation profiles of the age-DMRs shown in the browser view.

Using our model fits, we classified age-DMRs into four age-related dynamic patterns based on the direction and magnitude of changes in mCG and hmCG over time (Fig. 4D; S7D; Table S3): regions that show an increase in mCG and hmCG (Pattern 1); regions that show an increase in total mCG and hmCG, but a decrease in mCG (Pattern 2); regions that show a decrease in total mCG and mCG, but an increase in hmCG (Pattern 3); and regions that show a decrease in both mCG and hmCG (total demethylation pattern, Pattern 4) (Fig. 4G,H). Each pattern was further subdivided into ‘fast’ and ‘slow’ sub-patterns based on the mC oxidation time constant, τ_𝑚𝐶→ℎ𝑚𝐶_ .

The majority of DMRs progressively increased total mCG across the lifespan, falling into pattern 1 (56% of Glu and 75% of MGE-GABA age-DMRs) or pattern 2 (14% in both Glu and MGE-GABA). These patterns had significantly faster C-to-mC conversion ( τ_𝐶→𝑚𝐶_ < 1 year in fast subgroups) compared to patterns 3 and 4 (τ_𝐶→𝑚𝐶_ > 2 years). The C-to-mC transition rate was also faster than mC-to-hmC and hmC-to-C transitions within patterns 1 and 2 (Fig. 4D,E; S7F).

In contrast, pattern 3 age-DMRs (20% in Glu and 5% in MGE-GABA) had similar time constants for all three transitions (C to mC, mC to hmC, and hmC to C) (Fig. S7F). The small number of total demethylation age-DMRs (pattern 4, 4% in Glu and 3% in MGE-GABA) that showed higher hmCG at infancy, had much slower C-to-mC conversion rates: >4 years in the fast subgroup and >30 years in the slow subgroup. This slow conversion may indicate that the epigenetic state at these sites is linked to long-term gene activation or sustained chromatin accessibility. Age-DMRs in patterns 3 and 4 were enriched for DNA sequence motifs recognized by neuronal activity-regulated transcription factors (TFs), such as the AP-1 factors FOS and JUNB^62^, in both cell types (Fig. S7G). Interestingly, motifs for the activity-regulated TF EGR1 were enriched in age-DMRs in Glu neurons but not in MGE-GABA neurons.

To better understand the epigenetic context of age-DMRs, we used ChIP-seq to examine how age-related DNA methylation dynamics interact with histone modifications (Methods, Table S4–6). We observed the expected reciprocal relationship between DNA methylation and the active histone mark, H3K27ac, across development in both cell types (Fig. 4F). Age-increased H3K27ac regions were strongly enriched for age-DMRs in patterns 3 and 4, in which total mCG decreases with age. Conversely, age-decreased H3K27ac regions were enriched for age-DMRs in patterns 1 and 2, where total mCG increases with age. The only exception was the fast pattern 2 DMRs in MGE-GABA, which showed enrichment in both age-increased and age-decreased H3K27ac regions. Notably, in pattern 2 age-DMRs, gains in total mCG result mainly from increasing hmCG (Fig. 4D), suggesting that increased total mCG at these sites may be associated with either activation or deactivation of regulatory elements.

Age-DMRs in patterns 3 and 4, which undergo age-dependent demethylation, were depleted of all regions marked by the polycomb repressive complex 2-associated mark, H3K27me3. This suggests that active regulatory elements experiencing demethylation over postnatal development are largely distinct from Polycomb-repressed regions. In contrast, pattern 1 and 2 age-DMRs, which gain methylation with age, show more nuanced relationships with Polycomb regulation depending on genomic context (Fig. 4F). Pattern 1 age-DMRs enriched in age-increased H3K27me3 regions were preferentially localized to intergenic regions (Fig. S8A). By contrast, pattern 1 and 2 DMRs enriched in age-decreased H3K27me3 regions localized to CGIs and CGI shores (Fig. S8A). CGIs are found in ∼60% of promoters, and CGI shores play critical roles in gene regulation^63,64^. We previously identified a paradoxical pattern of high total mCG at CGI shores in promoters of several Glu-specific genes, including key transcription factors and major markers of Glu identity (e.g., *BDNF, EMX1, SLC17A6, PAX6, NRN1*)^37,64^. Here, we show that in these loci, progressive accumulation of hmCG from infancy to adulthood drives the emergence of high total mCG, replacing early-life Polycomb repression marked by H3K27me3 (Fig. S8B). Thus, pattern 1 and 2 age-DMRs that gain hmCG while losing H3K27me3 may be associated with genes that are actively expressed in adulthood despite age-related increases in total mCG.

Finally, we found that pattern 1 and 2 slow DMRs are strongly enriched in regions with age-decreasing heterochromatin (H3K9me3) in both Glu and MGE-GABA neurons (Fig. 4F). Many of these DMRs are within 1 Mb of centromeres across pericentromeric domains of all chromosomes (Fig. S8C,D), suggesting that loss of constitutive heterochromatin in these regions is accompanied by gradual accumulation of total mCG.

### Dynamic methylation of age-dependent enhancers

Active cis regulatory elements (CREs), including enhancers and promoters marked by H3K27ac, are critical drivers of cell-type-specific gene expression^65,66^. We identified candidate CREs marked by H3K27ac across all stages of postnatal brain development (159,442 unique regions in Glu, 145,758 in MGE-GABA neurons). The majority of CREs (77% in Glu, 79% in MGE-GABA) were distal (>1 kb) from the nearest transcription start site (TSS), representing putative enhancer regions. While ∼37% of CREs in Glu and ∼56% in MGE-GABA remained stable with age (constitutive CREs), a substantial proportion of CREs had significant age-related changes in H3K27ac ChIP-seq (99,699 age-differential peaks in Glu, 63,199 peaks in MGE-GABA; log2 fold-change > 0.5, FDR < 0.05) (Fig. 5A). Among age-differential CREs, >95% changed monotonically across the lifespan, with ∼40% showing progressive activation and ∼55% showing progressive deactivation. The remaining ∼5% had transient, non-monotonic patterns (4,279 in Glu and 469 in MGE-GABA) (Fig. 5A). The dense sampling of ages in our dataset allowed us to classify dynamic CREs based on the timing of changes in H3K27ac, distinguishing early (infancy to early childhood) versus late (early childhood to adulthood) activation or deactivation (Fig. 5A; Fig. S9A).

**Figure 5.**
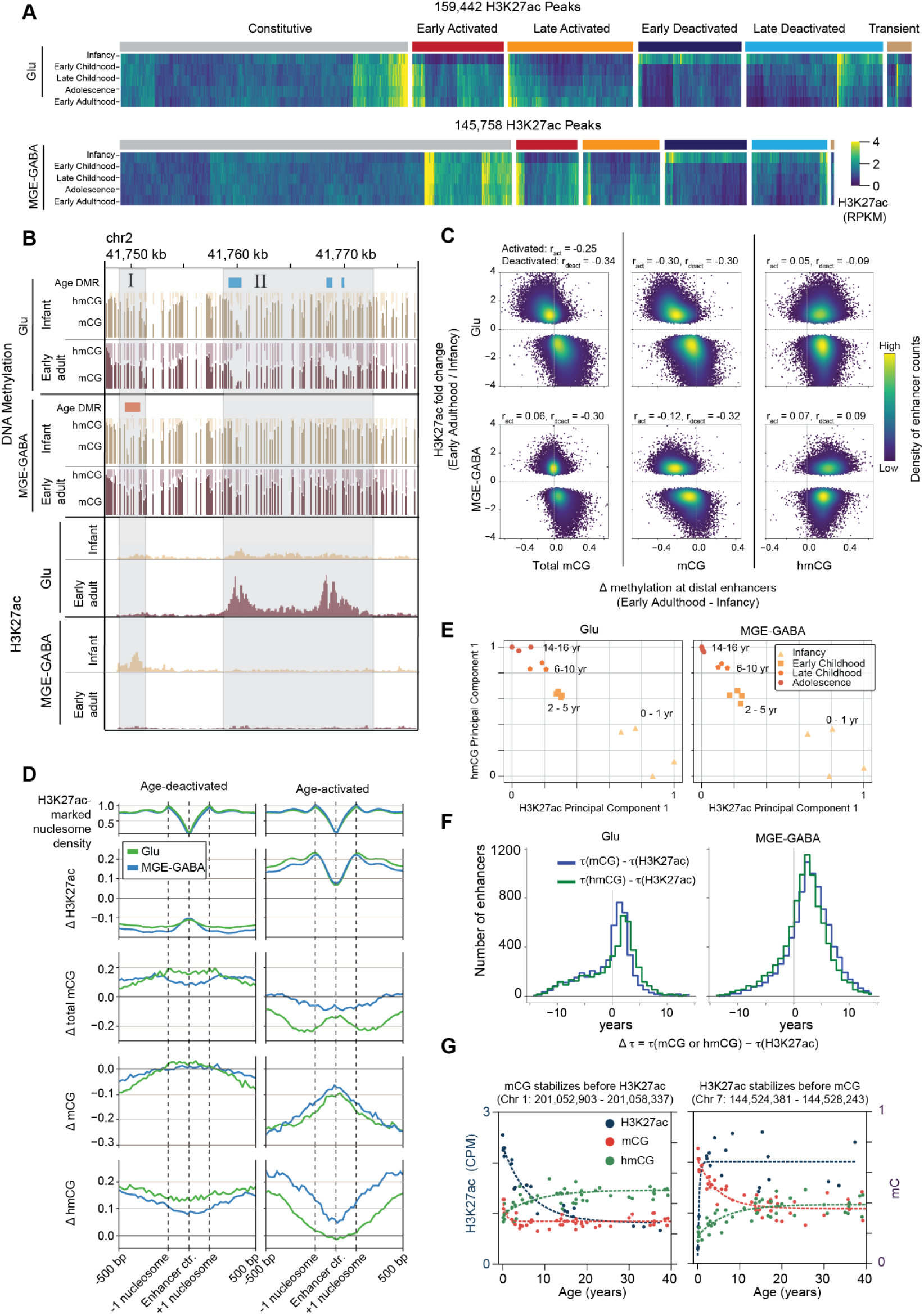
Coordinated regulation of enhancer activity via DNA methylation and H3K27ac. **(A)** Heatmaps illustrating the variation in age-related differential H3K27ac peaks across two cell types. Age differential peaks are classified as activated, deactivated, or transient, and further categorized as early- or late-changing. **(B)** Examples of cell-type specific enhancer regulation in an intergenic region near the gene *PKDCC*, showing coordinated changes in H3K27ac. I: An MGE-GABA enhancer in the infant brain that gains mC and loses H3K27ac. II: A Glu enhancer that loses mC and gains H3K27ac with age. **(C)** Scatter plots comparing developmental changes in DNA methylation with changes in H3K27ac for each differential peak identified when comparing early adulthood to infancy. **(D)** Histone modification and DNA methylation dynamics around precisely localized age-activated and age-deactivated enhancers, defined by open chromatin and H3K27ac peaks. Age-activated CREs gain hmCG in MGE-GABA, but not in Glu neurons, highlighting cell type-specific epigenetic regulation. Density of H3K27ac-marked nucleosomes is estimated from MNase-digestion-based ChIP-Seq data. **(E)** Principal component analysis shows coordinated, synchronic developmental trajectories of H3K27ac and hmCG. **(F)** Divergent timing of enhancer activation and DNA methylation changes at individual CREs. **(G)** Examples of CREs in Glu neurons with rapid mCG and gradual H3K27ac dynamics (left), or rapid H3K27ac and gradual mCG (right).

To investigate the functional role of distal CREs (putative enhancers, Table S4), we linked each enhancer with the nearest TSS and analyzed functional enrichment of genes that were expressed in our RNA-seq dataset (Fig. S9B, Table S7). In Glu neurons, age-activated enhancers (both early and late) were located near genes involved in synaptic signaling, trans-synaptic communication, and neuronal maturation, consistent with the postnatal activation of excitatory neuronal function (FDR < 1e-3). Age-deactivated enhancers, by contrast, were associated with genes involved in early neurodevelopmental processes, such as neurogenesis, cell morphogenesis, and neuron projection development. Constitutive enhancers were broadly linked to genes involved in core cellular processes like cell signaling, suggesting a role in maintaining baseline Glu neuron identity. We observed similar enhancer–gene relationships in MGE-GABA neurons (Fig. S9B), indicating that these dynamic regulatory principles are conserved across major neuronal subtypes.

Although total mCG is inversely associated with enhancer activity during development^67^, the relative contribution of mCG versus hmCG has not been defined. Our data illustrate the complex relationship among these epigenetic marks, with examples of opposite cell type-specific developmental regulation at two neighboring CREs in an intergenic region (Fig. 5B). At these sites, developmental changes in H3K27ac are inversely correlated with changes in total mCG. We analyzed the correlation between H3K27ac and DNA methylation separately for age-activated and -deactivated enhancers. Age-deactivated enhancers had a strong negative correlation with total mCG in both neuron types (Fig. 5C). In contrast, among age-activated enhancers, this relationship was observed in Glu but not in MGE-GABA cells, highlighting a distinct methylation-activity relationship in MGE-GABA neurons (Fig. 5C).

To assess the coordinated relationship between H3K27ac and DNA methylation at age-activated and age-deactivated enhancers in more detail, we estimated enhancer centers defined by H3K27ac (Fig. S9C). We also developed a more precise method for localizing enhancers by estimating nucleosome positions using our H3K27ac MNase-ChIP-seq data employing DANPOS^68^, and combined these with open chromatin regions from single-nucleus ATAC-seq^69^ (Fig. 5D, S9D; Table S8, Methods). Age-deactivated enhancers had increasing total mCG in both Glu and MGE-GABA neurons (Fig. 5D, Fig. S9C, I-II). Notably, this increase was driven primarily by gains in hmCG, with minimal net changes in mCG (Fig. 5D, Fig. S9C, III-VI). This suggests that in age-deactivated enhancers, de novo DNA methylation was followed by subsequent conversion of mCG to hmCG.

In contrast, age-activated enhancers had distinct DNA methylation dynamics in Glu versus MGE-GABA neurons. These enhancers in Glu neurons underwent robust DNA demethylation (loss of total mCG) (Fig. S9C, VII). MGE-GABA neurons did not show this pattern, and instead had largely conserved total DNA methylation at age-activated enhancers. This is consistent with a balance between mCG loss and hmCG gain in MGE-GABA cells (Fig. S9C, VIII). Using our high-precision estimates of enhancer centers, we found that this difference between Glu and MGE-GABA neurons occurs primarily in the flanks of the open chromatin regions, corresponding to the neighboring nucleosomes (Fig. 5D).

To understand the regulatory networks that could shape cell type-specific epigenetic dynamics, we analyzed the enrichment of DNA sequence motifs associated with TF binding within ±50 bp of the age-dependent enhancers^62^ (Fig. S9E, Table S9). Age-activated Glu enhancers showed a strong cell type-specific enrichment for the binding motif of EGR1, a key activity-dependent TF that interacts with TET1 in mouse cortex^70^. EGR1 is most strongly expressed in Glu neurons, and peaks around age 2 coinciding with a critical stage of circuit refinement (Fig. S9F). We propose that in Glu age-activated enhancers, EGR1 binding may recruit TET1 to promote hmCG turnover and demethylation at adjacent H3K27ac-marked nucleosomes. The lack of EGR1 motif enrichment in MGE-GABA age-activated enhancers suggests this mechanism may be specific to excitatory neurons.

We took advantage of our paired measurements of DNA methylation and H3K27ac in the same tissue samples to explore the relative timing of changes in these two epigenetic marks during development. When comparing the overall pattern of change using the first principal components of H3K27ac and of hmCG (Fig. 5E) or mCG (Fig. S9G), we found that H3K27ac changed in parallel with mCG and hmCG, with no major difference in their overall timing. We further investigated the timing of dynamic regulation at individual enhancers. For each age-dependent enhancer, we fitted exponential models to the time course of H3K27ac, mCG, and hmCG. We focused on regions with high-quality fits across all three modalities (R² ≥ 0.4; 4,667 regions in Glu, 8,842 in MGE-GABA, Fig. 5F). At most enhancers, H3K27ac changes preceded DNA methylation changes (55.8% of enhancers in Glu, 81.4% in MGE-GABA). However, there was a broad distribution of time scales, with differences between the two modalities as large as ∼10 years at specific regions. We illustrate the extremes of this distribution with two examples of distal enhancer regions in Glu neurons that lose mCG in early infancy followed by more gradual loss of H3K27ac (Fig. 5G, left), or that rapidly gain H3K27ac in infancy followed by a gradual loss of mCG (Fig. 5G, right).

### Epigenetic regulation of age-related gene expression in neurons

In the human brain, DNA methylation at CG and non-CG sites is strongly correlated with cell type specific gene expression in adult neurons^5,8,36–38,71–73^, but the relationship of dynamic (hydroxy)methylation changes with age-dependent gene expression is poorly characterized^17,29,74^. We performed RNA-seq from FANS-isolated Glu and MGE-GABA neuronal nuclei, identifying 8,109 unique age-related differentially expressed genes (age-DEGs, FDR < 0.05; Fig. S10A; Table S10). We found more age-DEGs in Glu (6,808) than in MGE-GABA neurons (4,929), potentially reflecting the greater cellular diversity of the Glu population. Notably, 43.2% of age-DEGs were shared between Glu and MGE-GABA neurons, with 95.5% of these showing consistent directionality (Fig. S10B). We found 157 genes with opposite patterns of age-related regulation in the two cell types, including the proline transporter *SLC6A7*, *ADGRG6*, and *EXPH5* (Fig. 1E; S10B). *ADGRG6* encodes an adhesion G protein–coupled receptor implicated in developmental signaling and neuronal maturation, whereas *EXPH5* encodes a Rab27 effector implicated in intracellular vesicle trafficking^75,76^. Across all comparisons, downregulated genes outnumbered upregulated genes by ∼30%, consistent with cortical snRNA-seq data from developmental and aging studies^29,77,78^ (Fig. S10C-E).

We classified age-DEGs in Glu and MGE-GABA neurons into five dynamic patterns (Early-Up, Early-Down, Late-Up, Late-Down, and non-monotonic), capturing expression changes from infancy through adulthood (Fig. 6A, B; Table S11). Following a marked shift from infancy to early childhood, age-DEGs continued to accumulate, though to a lesser extent, through childhood to adolescence. Only a small subset of genes (497 in Glu, 50 in MGE-GABA) had significant changes in expression after adolescence. Functional enrichment analysis revealed sustained downregulation from infancy to adolescence of pathways related to synaptic organization, neurite development, cell adhesion, protein synthesis, and ion transport, consistent with early neurodevelopmental processes such as synaptic pruning (Fig. S10F; Table S12). In contrast, upregulated genes had fewer functional enrichments, including ECM remodeling and serine hydrolase activity (Fig. S10G; Table S12). These findings are broadly consistent with recent snRNA-seq analyses of neuronal development in the human DLPFC^79–81^.

**Figure 6.**
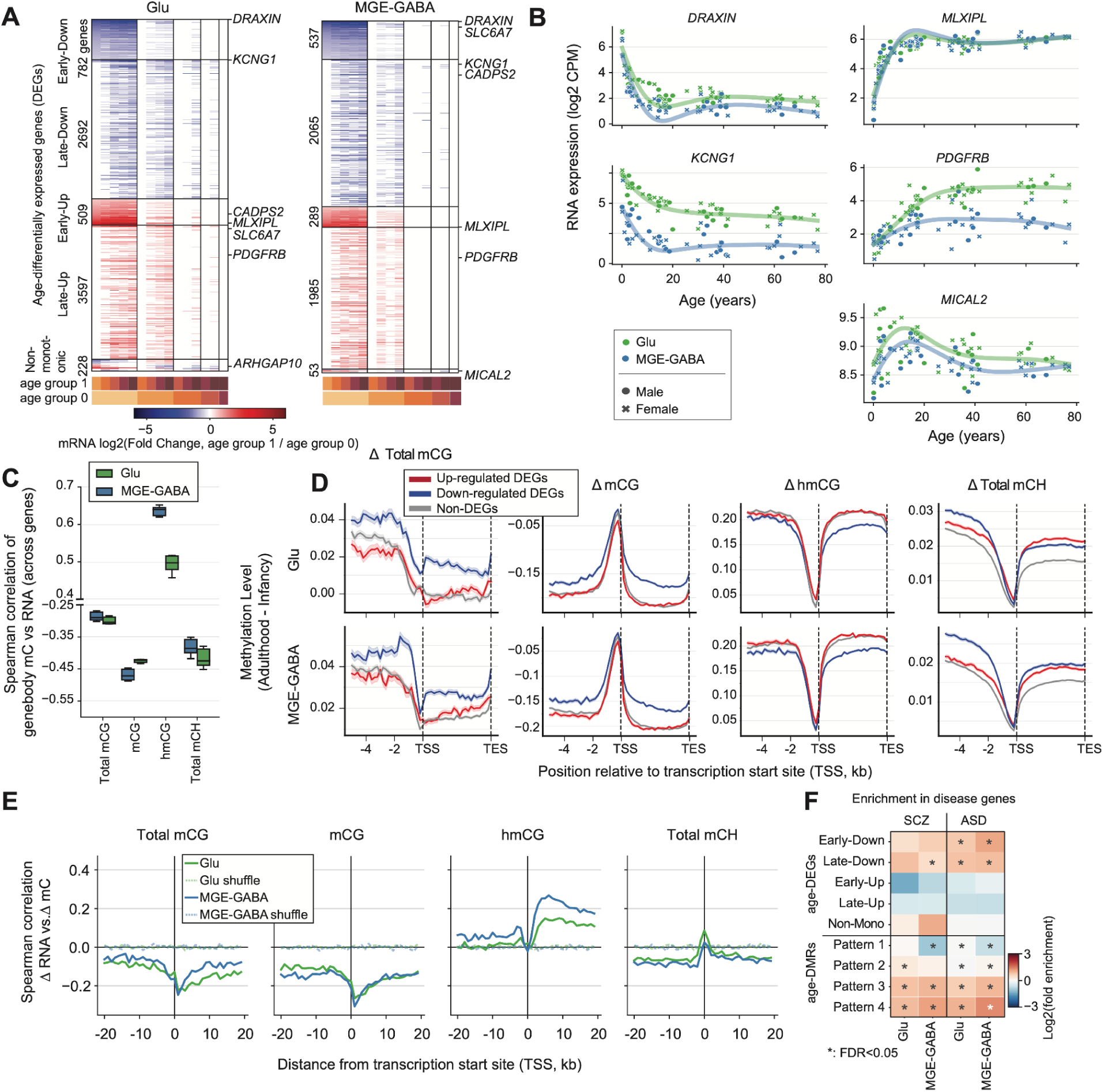
Age-dependent gene expression dynamics and epigenetic regulation. **(A)** Heatmap of age-DEGs clustered into five groups based on expression patterns: early downregulated (Early-Down), late downregulated (Late-Down), early upregulated (Early-Up), late upregulated (Late-Up), and non-monotonic (Non-Mono). **(B)** Age-dependent RNA expression (log2CPM) of selected genes from each group. **(C)** Spearman correlation between RNA expression gene body methylation across all genes within each period. **(D)** Methylation changes between adulthood and infancy within gene bodies and 5kb upstream regions for upregulated, downregulated, and non-differentially expressed genes. **(E)** Spearman correlation between RNA expression fold change and DNA methylation fold change within the flanking regions of TSS of age-DEGs across all pairwise comparisons (solid line). Dashed lines represent the correlation between shuffled fold changes. **(F,G)** Enrichment of schizophrenia- and ASD-associated genes among age-DEGs (F) and age-DMRs (G).

We confirmed that, in each age group, gene expression has a negative correlation with total mCG, mCG and mCH, and a positive association with hmCG in the gene body^16^ (Fig. 6C, Spearman correlation |r| = 0.18-0.65). To connect age-related epigenetic dynamics with gene expression, we assessed DNA methylation changes from infancy to adulthood across upregulated, downregulated, and stably expressed genes (Fig. 6D). We found that age-upregulated and stably expressed genes experience a greater conversion of mCG to hmCG within their gene body compared with age-downregulated genes. In age-downregulated genes there was a net increase in total CG methylation (mCG+hmCG), consistent with less TET-mediated demethylation. These differences were not explained by the initial expression level in infancy, suggesting they reflect dynamic post-natal regulation of transcription (Fig. S11A, Mann-Whitney U test, p < 0.05).

To further characterize the relationship of dynamic RNA expression with DNA methylation, we correlated in 1kb bins flanking the TSS of age-DEGs across all pairs of age groups. Within gene bodies, mCG changes were strongly negatively correlated with expression changes, whereas hmCG changes were positively correlated (Fig. 6E). These findings suggest divergent functional roles, with mCG repressing and hmCG activating gene expression through the lifespan.

In contrast with mCG, we found that mCH had a more complex, non-monotonic relationship with age-related gene expression. Both up- and downregulated genes accumulated more mCH than non-age-differentially expressed genes (p<0.05, Fig. S11A). We previously reported that genes with cell type-specific expression (e.g. Glu > MGE-GABA or MGE-GABA > Glu) have higher gene body mCH than non-cell-type-specific genes with the similar expression level^37^. Consistent with this, our data indicate that elevated gene-body mCH marks genes that are dynamically regulated with age, regardless of direction. Gene body mCH changes with age had a strong negative correlation with gene expression changes from adolescence to adulthood (Fig. S11C), consistent with previous findings linking mCH to gene repression in the adult brain^5,29^. By contrast, during early development, when mCH accumulates genome-wide, we observed little association between mCH gain and gene expression changes between infancy and adolescence (Fig. S11B). Thus, while early-life mCH accumulation may not directly correlate with gene expression, mCH gains later in life are more closely tied to transcriptional repression.

Our data present an opportunity to functionally interpret the genetic sequences associated with risk for neurodevelopmental disorders. To link the developmental dynamics of Glu and MGE-GABA neurons with their potential impact on disease, we compared our age-DEGs and age-DMRs with large-scale genome-wide association studies for neurodevelopmental disorders. Genes associated with schizophrenia (SCZ)^82^ were enriched in late-downregulated age-DEGs in MGE-GABA neurons (Fig. 6F). Genes associated with autism spectrum disorder (ASD)^83^ were enriched in both early- and late-downregulated genes in both MGE-GABA and Glu neurons (Fig. 6F). We found that Patterns 3 and 4 age-DMRs, which decrease total methylation during development, were especially enriched for genes associated with SCZ and ASD (Fig. 6G).

## Discussion

The discovery that hydroxymethylcytosine (hmC) is highly enriched in adult mammalian neurons^11^ and generated from mC via TET dioxygenase enzymes^10^ raises the question of what functional role this abundant neuronal epigenetic modification plays in brain development and function. The disruption of neurodevelopment and cognition in Rett’s syndrome, caused by loss of function in the DNA methylation reader MECP2, highlights the critical importance of DNA methylation in neuronal maturation^21^. The recent recognition of Beck-Fahrner syndrome^22^, a neurodevelopmental disorder caused by mutations disrupting the DNA methylcytosine dioxygenase TET3, further suggests that developmentally patterned hydroxymethylation is essential for healthy neuronal function. We comprehensively profiled the genome-wide lifespan dynamics of mC and hmC in human excitatory and inhibitory neurons, explicitly resolving their interconversion rather than total methylation alone. Our dataset, including post-mortem brain samples from 103 donors, reveals coordinated cell-type specific development of mC and hmC in parallel with transcriptional and chromatin changes. Our findings position hmC as a major component of neuronal DNA methylome dynamics with broad implications for epigenetic regulation of brain function.

Two major hypotheses have proposed that hmC may be a transient epigenetic modification acting as an intermediate in active DNA demethylation, or else that hmC is a stable component of the methylome with a direct role in modifying gene regulation^12,17,25,37,84^. Our data showed that neurons experience a dramatic conversion of mCG to hmCG that unfolds over the first two decades of life, coinciding with the stabilization of neuronal identity and synaptic architecture. The conversion of mCG to hmCG ultimately affects around half of all CG sites in the genome. The scale, cell-type specificity, and persistence of this shift over decades are inconsistent with hmC acting solely as a transient demethylation intermediate, which would predict rapid turnover rather than long-term accumulation. This transformation parallels the genome-wide accumulation of mCH in neurons^5^, which occurs during the same period of neuronal maturation and synaptic proliferation, refinement and pruning. Together, mCH and hmCG represent unique neuron-specific forms of DNA methylation that profoundly shape the epigenetic landscape during postnatal brain development.

We found that a simple, three-state dynamical model for the cyclic interconversion of C, mC and hmC captures the complex trajectory of DNA methylation across the lifespan. A greater proportion of mCG converted to hmCG in Glu compared to MGE-GABA neurons^36,37^, suggesting that the distinct functions of excitatory and inhibitory neurons require specialized epigenetic regulation. Future studies with single-cell resolution will help to refine our understanding of mC and hmC dynamics in subtypes of glutamatergic and GABAergic neurons^33,34,39^. Notably, cell type- and age-dependent gene expression patterns of methylcytosine dioxygenases (TETs) and DNA methyltransferases do not fully explain the dynamics of mC and hmC. Instead, these dynamics likely reflect additional layers of regulation, such as the dependence of TET enzyme activity on availability of essential cofactors (e.g., Fe²⁺) and co-substrates (α-ketoglutarate)^10,85^. This hypothesis suggests a testable model in which neuronal metabolic state—via availability of Fe²⁺ and α-ketoglutarate—modulates TET activity and thereby sculpts cytosine modification landscapes during development.

Our findings complement research on DNA methylation-based biological clocks, i.e. statistical models which accurately predict the age of an individual based on average methylation patterns at several hundred CG sites^4,47,48,50,51^. We found a clock-like, linear increase of the strand asymmetry of hmC within the body of highly expressed genes across the entire lifespan^36,46^. This clock signal is strongly correlated with age (Pearson |r| = 0.97-0.98), on par with or even more accurate than previous DNA methylation-based clocks (r = 0.935-0.991 for brain tissue^51^). Unlike prior methylation clocks, the average hmC strand asymmetry correlates directly with age without any need for fitting a regression model with hundreds of gene-specific weight parameters. The robust, linear relationship between strand asymmetry, gene expression, and age is consistent with a mechanistic model in which the association of transcription machinery with DNA on the template (antisense) strand inhibits conversion of mC to hmC.

Previous studies have shown that gene-body hmC is positively associated with gene expression^12,16^. Here, we extend this relationship to developmental dynamics: for a given gene, age-dependent changes in expression are positively correlated with changes in hmC^86^. Moreover, the stronger correlation in gene bodies compared to promoter regions suggests that gene body methylation may be more tightly linked to age-related transcriptional changes. These findings are consistent with a model in which hydroxymethylation is permissive for gene expression.

We identified thousands of age-DMRs across the genome in Glu and MGE-GABA neurons during postnatal development, revealing extensive locus-specific remodeling of total CG methylation (mC+hmC) and resolving the underlying contributions of mCG and hmCG dynamics. Many age-DMRs were cell-type-specific, including some with opposite age-dependent methylation trends in Glu and MGE-GABA neurons. These findings substantially expand the catalog of neuronal age-DMRs^5,29,43^ and underscore the extensive epigenomic remodeling that accompanies neuronal maturation across the postnatal lifespan.

Age-related DNA methylation dynamics are bidirectionally coordinated with changes in histone modifications. Regions that lose methylation are strongly enriched for age-increased H3K27ac, consistent with activation of promoters and enhancers during neuronal maturation. Developmentally activated enhancers in Glu neurons were enriched in the binding motif for EGR1, suggesting this neuronal activity-related TF may recruit TET1 to promote demethylation of enhancers specifically in excitatory but not inhibitory neurons^70^. Although the average genome-wide pattern of H3K27ac changed in parallel with mC and hmC during development, we identified specific regions where histone modification changes preceded DNA methylation or vice versa. These findings align with data from neural progenitor cells showing temporally discordant waves of open chromatin and DNA methylation changes at thousands of genomic loci^87^.

Our data link developmental increases in total DNA methylation with loss of Polycomb repressive complex 2, marked by H3K27me3, in CGIs and CGI shores. These changes point to a shift from Polycomb repression to active transcription despite high DNA methylation. This finding challenges the view that DNA methylation is inherently repressive. Instead, hydroxymethylation may support sustained gene expression even in regions with high total methylation. Hydroxymethylation may therefore stabilize transcriptional programs after developmental repression is removed, helping maintain neuronal identity over time. We also found a coordinated relationship between mCG dynamics and the constitutive heterochromatin mark H3K9me3 at pericentromeres^86^.

Our findings resolve the specific forms of DNA methylation at CG and non-CG sites which contribute to female-specific X chromosome inactivation^5,55^. We found that the inactive X chromosome (Xi) has a net loss of total CG methylation (mCG+hmCG) during neuronal development driven primarily by decreased hmCG. By contrast, the average level of mCG on Xi remains largely stable. Promoter hypermethylation at inactivated genes in females is due to mCG rather than hmCG, reinforcing the role of promoter mCG in maintaining gene silencing^37,88^. Together, these findings suggest that heterochromatin regions such as Xi are inaccessible to TET-mediated oxidative conversion of mCG to hmCG, reinforcing the silenced state of inactivated genes and maintaining dosage compensation^89^.

In conclusion, our study comprehensively characterizes the complex and dynamic regulation of DNA methylation and hydroxymethylation in human neurons across the lifespan. The distinct patterns of age-related changes in Glu and MGE-GABA neurons underscore the importance of cell-type-specific epigenetic regulation in maintaining neuronal development and aging. Our findings of the distinct dynamical roles of mC and hmC have important implications for future research on neurodevelopmental and neurodegenerative diseases, and for developing potential therapeutic strategies targeting epigenetic modifications. More broadly, these results motivate mechanism-aware biomarkers and therapeutic approaches that distinguish methylcytosine states rather than treating DNA methylation as a single entity.

## Data availability

Raw data from this study have been deposited at the NIMH Data Archive (NDA) in collection 5862. An interactive data browser for visualization and analysis is available at https://brainome.ucsd.edu/BrainHydroxyDevelopment.

## Acknowledgments

We are grateful to M.M. Behrens, J.R. Ecker, and C. Luo for helpful discussions. This project was supported by NIH/NIMH grants U01MH122590 (SD), U01MH122591 (CL) and U01MH122592 (EAM), and by VA Career Scientist Award K6BX006524 (SD).

## Methods

### Human brain samples

Human postmortem brain tissues were sourced from the dorsolateral prefrontal cortex (Brodmann area 9). The majority of the samples were obtained from the NIH NeuroBioBank. Twenty nine adult samples were obtained from the New York State Psychiatric Institute Macedonian Brain Collection. Specific characteristics of the individuals are displayed in Table S1. The dataset included 14 donors diagnosed with schizophrenia (SCZ), ranging in age from 20 to 60 years. No clear separation between SCZ and non-SCZ donors was observed in the top two principal components of DNA methylation or gene expression (Fig. 1C). Similarly, no significant differences were detected in genome-wide methylation levels or in the time constants for methylation conversion (Fig. S1E,F). Variance partitioning further indicated minimal disease-associated effects (Fig. S1G). Nevertheless, SCZ samples were excluded from further, except where noted to avoid potential confounding effects.

### Nuclei isolation

The protocol and reagents for brain nuclei isolation before flow cytometry were previously described^36,37^. Briefly, tissue was homogenized in a lysis buffer, underlaid with a high sucrose buffer, and centrifuged for 1 hour. The nuclear pellets were resuspended in an antibody-incubation buffer, incubated with primary antibodies for 1.5 hours, followed by a second centrifugation and a 1-hour incubation with secondary antibodies. We used FANS, employing NeuN (RBFOX3) as a marker for neuronal nuclei and SOX6 as a marker of MGE-derived GABA neurons, to separate nuclei of MGE-GABA and Glu neurons as NeuN+/SOX6+ and NeuN+/SOX6- populations, respectively. The following antibodies were used for FANS: (1) mouse monoclonal anti-NeuN phycoerythrin (PE)–conjugated antibodies (Millipore, FCMAB317PE, 1:1000), (2) guinea pig polyclonal anti-SOX6 antibodies^90^, 1:1500 (3) donkey anti-guinea pig AX647-conjugated secondary antibodies (Jackson ImmunoResearch, 1:1500).

### RNA isolation and RNA-seq library construction

For RNA isolation, nuclei were prepared using the FANS-based method as described above, with the addition of RNase inhibitors (Recombinant RNAse Inhibitor, Takara Bio; 1:40 dilution during lysis step, 1:100 during other steps). ∼40,000 sorted nuclei per sample were used for RNA extraction with the PicoPure RNA Isolation Kit (Thermo Fisher). RNA-seq libraries were prepared from 10 ng of RNA using the SMARTer® Stranded Total RNA-Seq Pico Input Mammalian Kit v2 SMARTer Stranded Total RNA-Seq Kit (Takara Bio) and sequenced on a HiSeq 2500 with the paired-end 100 cycles protocol, to an average of ∼100 million read pairs per sample.

### DNA methylation assay

Between 350,000 and 700,000 nuclei of Glu and MGE-GABA neurons were isolated from each tissue sample by flow cytometry. High-quality genomic DNA (average fragment length >20,000 bp) was isolated using the DNeasy Blood and Tissue Kit (Qiagen) and concentrated with a Genomic DNA Cleanup Kit (Zymo). Genomic DNA (500 ng) was used for BS/OxBS processing and library preparation with the TrueMethyl Whole Genome kit (CEGX). Libraries were sequenced on an Illumina HiSeq 2500 system using a paired-end 100-cycle protocol. Conversion performance in oxidation and bisulfite treatments was assessed using CEGX spike-in control oligonucleotides.

### Histone modification ChIP-seq

For each ChIP-seq experiment, between 100,000 and 150,000 nuclei of Glu and MGE-GABA neurons were isolated from each tissue sample by FANS, as described above and in previous work^37^. During the protocol, lysis buffer and immunoprecipitation (IP) buffer were supplemented with protease inhibitors 0.1 mM benzamidine and 0.1 mM phenylmethylsulfonyl fluoride (PMSF). For ChIP-seq, we followed a previously optimized and extensively tested native ChIP protocol (N-ChIP) in which chromatin is fragmented into individual nucleosomes using micrococcal nuclease (MNase) without crosslinking proteins to DNA^91^. For the IP reactions, we used the following commercial ChIP-grade antibodies against histone modifications: (1) anti-H3K27ac antibodies from Active Motif (Cat# 39133; rabbit polyclonal, 3 µg per sample), (2) anti-H3K27me3 antibodies from EMD Millipore (Cat# 07-449; rabbit polyclonal, 3 µg per sample), (3) anti-H3K9me3 antibodies from Abcam (Cat# 8898; rabbit polyclonal, 3 µg per sample). ChIP-Seq libraries were prepared with the NEBNext Ultra II DNA Library Prep Kit for Illumina (New England Biolabs, Cat. # E7645). The resulting libraries were sequenced on a NovaSeq 6000 instrument (Illumina), using paired-end 100 cycles protocol, to an average of 60 million read pairs per sample. For each cell type and age group, two input control samples obtained from MNAse-digested DNA were prepared and sequenced.

### RNA-seq data processing and mapping

RNA reads were trimmed using Trim-Glalore (0.6.4_dev) for quality trimming and adaptor trimming. The first 3 bps from the 5’ end of read1 is further removed. Trimmed reads were then mapped to hg38 and GENCODE gene annotation (v37) using STAR (v2.7.3a) with the following parameters consistent with ENCODE standard ( --outFilterType BySJout --outFilterMultimapNmax 20 --outFilterMismatchNmax 999 --outFilterMismatchNoverReadLmax 0.04 --alignIntronMin 20 --alignIntronMax 1000000 --alignMatesGapMax 1000000 --alignSJoverhangMin 8 --alignSJDBoverhangMin 1 --sjdbScore 1). Gene count was estimated by RSEM(v1.3.1) with the following parameters (--estimate-rspd --paired-end --forward-prob 1).

Samples were excluded from downstream analysis if they exceeded a 5% mitochondrial read ratio or showed abnormal gene expression patterns based on correlation and PCA with other samples.

### Mapping DNA methylation data

Whole genome (oxidative)bisulfite sequence was mapped to hg38, C to T converted genome index in paired-end by Hisat-3N (v2.2.1-3n-0.0.3) after adaptor trimming by trim_galore (v0.6.4_dev). Only unique mapped reads were kept for further analysis. Duplicated reads were removed by Picard (v2.27.1). The number of methylated and unmethylated reads covered for each cytosine site was called by methylpy (v1.4.4).

Samples were excluded from downstream analysis if they had an average cytosine coverage below 5x, lacked clear separation between cell types, or exhibited abnormal non-CG methylation patterns. For each donor, cytosines and their adjacent 3’ sites overlapping with homozygous single nucleotide polymorphisms (SNPs) identified in any of the donors were excluded from downstream methylation analysis.

### Mapping Histone modification ChIP-seq data

Raw sequencing data were pre-processed to remove adapters and low-quality sequences with the HTStream tool (https://github.com/s4hts/HTStream). Reads were mapped to the human hg38 reference genome with BWA-MEM^92^, and filtered to remove multi-mapping reads and low-quality alignments using Samtools^92^. Reads mapping to ENCODE-blacklisted genomic regions were excluded using BEDTools^93^,

### First-order dynamical model

We fit a first-order exponential model of methylation with age, 𝑦_𝑚𝑒𝑡ℎ𝑦𝑙𝑎𝑡𝑖𝑜𝑛_ = 𝑎 · 𝑒𝑥𝑝 (− 1/τ · 𝑥_𝑎𝑔𝑒_) + 𝑐 , using scipy.optimize.curve_fit to estimate the parameter τ and its standard error.

### Differential expression and gene ontology

Protein coding and lncRNA genes from chromosomes 1-22 and X with counts per million (CPM) >7 in at least 3 samples are defined as “expressed genes”. Age differential expression testing was performed separately for each cell type and each comparison using Dream^94^ (variancePartition v1.28.9, edgeR v3.40.2). Specifically, calcNormFactors was used to control for the raw library size from the raw counts. Then, voomWithDreamWeights and dream were used to prepare for the linear mixed model fit from the normalized count and design matrix. The model included fixed effects of age group, batch, PMI hours, RIN,race and sex: 𝑒𝑥𝑝𝑟𝑒𝑠𝑠𝑖𝑜𝑛 ∼ 𝐴𝑔𝑒 𝐺𝑟𝑜𝑢𝑝 + 𝑏𝑎𝑡𝑐ℎ + 𝑃𝑀𝐼 ℎ𝑜𝑢𝑟𝑠 + 𝑅𝐼𝑁 + 𝑅𝑎𝑐𝑒 + 𝑆𝑒𝑥. Finally, after testing the generalized linear fixed model for each gene, genes with an adjusted p-value < 0.05 in any pairwise age group comparison are defined as age-related differentially expressed genes (age-DEGs) in our analysis. Only donors without Schizophrenia were included in the DEG analysis.

For genes exhibiting monotonic changes, we compared the fold change between early childhood and infancy to the fold change between adulthood and early childhood. Those genes are resulted in four categories: early downregulated genes, late downregulated genes, early upregulated genes, and late upregulated genes. Additionally, we identified genes showing non-monotonic change patterns.

Functional gene ontology (GO) enrichment analysis was performed using clusterProfiler (v 3.14.3)^95^. All expressed genes in each cell type were included as the background gene list. Gene ontology with gene size between 25 and 500 were included in the enrichment test. The Benjamini-Hochberg method was used to control the false discovery rate. Results from biological pathway (BP) and molecular function (MF) are shown.

### Differential methylation analysis

Differentially methylated regions (DMRs) were detected for each cell type and pairwise age group using DSS^96,97^ (DSS v2.34.0, bsseq v1.22.0). DMLfit.multiFactor was applied with model formula: 𝑚𝐶𝐺∼𝑎𝑔𝑒 𝑔𝑟𝑜𝑢𝑝 + 𝑠𝑒𝑥, with a smoothing window size 500 bp. Next, DMLtest.multiFactor was used to call differentially methylated loci. Finally, callDMR was used to merge significantly differentially methylated loci (p < 0.01) with a distance smaller than 50 bp. Merged regions with a length greater than 50 bp and containing at least 3 CpG sites are called differentially methylated regions (DMRs). Only donors without Schizophrenia were included in the DMRs analysis. DMRs with score (areaStat) > 15 were reported. To create a comprehensive set of DMRs, results from all age comparisons were merged. The resulting merged DMRs were then filtered to include only those with a mean total mCG level greater than 0.3 and a mCG range (maximum - minimum) greater than 0.3. These filtered DMRs were used as input for the three-state DNA methylation kinetic model (see Methods: DNA methylation 3-state kinetic model).

### Peaks Calling and Differential Peaks

To call peaks from the H3K27ac, H3K27me3, and H3K9me3 histone modification data, we used different tools and parameters optimized for each mark. For H3K27ac, we applied *MACS2*^98^ in narrow peaks mode with a threshold of -q 0.001, calling peaks for each individual sample. For H3K27me3, we used *Epic2*^99^, with parameters –bin-size 200 and –gaps-allowed 3. For H3K9me3, we applied *MACS2* in broad peak mode with parameters --broad-cutoff 0.01 -q 0.01. After calling peaks from each sample, for each ChIP-seq dataset within a given cell type, we aggregated peaks that were present in at least three samples across ages, merged overlapping peaks, and removed those overlapping blacklist regions.

To identify age-differential peaks for each histone modification and cell type, we first used csaw^100^ to calculated the normalization factor using high-abundance windows(1kb for H3K27ac and 10kb for H3K27me3 and H3K9me3). We then applied edgeR^101^ to test the significant age differential peaks using pairwise comparison between age groups. Peaks were defined as age-differential if they were significant in at least one comparison with FDR < 0.05 and |log2(fold change)| ≥ 0.5. To characterize the dynamics pattern among these age-differential peaks, we classified a peak as monotonic if all significant pairwise comparisons showed changes in the same change direction, and as transient otherwise. Within the monotonic group, we further distinguished early- versus late-changing peaks by comparing fold-change magnitudes between infancy and early childhood versus early childhood and early adulthood.

### Refinement of enhancer positions by integrating H3K27ac and ATAC-seq data

Active enhancers are characterized by TSS-distal open chromatin regions (OCRs) that harbor TF binding sites and are flanked by H3K27ac-marked nucleosomes^102^. To link the H3K27ac-marked regions from our study to functionally relevant OCRs and putative TF binding sites, we constructed a refined set of regulatory elements by integrating our nucleosome-resolved H3K27ac ChIP-seq profiles and single-nucleus ATAC-seq data from postnatal (neonatal-to-adult) DLPFC brain samples^69^.

First, from the single-nucleus ATAC-seq study (NIH GEO accession no. GSE204684), we obtained the OCR peak positions, as well as the ATAC-seq signal traces aggregated across postnatal stages (neonatal, childhood, adolescence, and adult) for Glu and MGE-GABA neurons. In both neuronal cell types, for each OCR we then calculated the summit, defined as the 1 bp position of maximum ATAC-seq signal, using a custom R script. These OCR summits were subsequently filtered by overlapping them with our Glu and MGE-GABA H3K27ac ChIP-seq peaks, allowing for a ±150 bp flanking window, which approximates the span of one nucleosome.

Next, we required that the H3K27ac peak-overlapping OCR summits fall within nucleosome-free regions flanked by H3K27ac-marked nucleosomes in Glu and MGE-GABA neurons. Our MNase-digestion–based ChIP-seq protocol inherently produces nucleosome-sized fragments, providing nucleosome-level resolution of H3K27ac profiles^91^. Using these data, we applied DANPOS^68^ (“danpos.py dpos -m 1 -p 1e-8 -q 0”, https://github.com/sklasfeld/DANPOS3) to identify the positions of H3K27ac-positive nucleosomes in each cell type. For this analysis, infant and adult H3K27ac datasets were each subsampled to 70 million aligned reads (using samtools and a custom script) prior to merging into the combined H3K27ac consensus BAM files for each cell type. The OCR summits were then compared with the DANPOS-called nucleosome positions. We retained only those OCR summits that ***(i)*** lay within an H3K27ac nucleosome-free region and ***(ii)*** were flanked within ±150 bp by H3K27ac-positive nucleosomes.

The resulting sets of autosome OCR summits (60,443 for Glu and 45,893 for MGE-GABA) represent the centers of ‘refined’ regulatory elements, including both promoters and enhancers. To classify the refined enhancers by developmental dynamics, we overlapped them with age-dynamic distal H3K27ac peaks. This analysis identified 13,435 age-upregulated and 13,985 age-downregulated refined enhancers in Glu neurons, and 6,916 age-upregulated and 7,138 age-downregulated refined enhancers in MGE-GABA neurons (Table S8). For TF binding prediction with HOMER^62^, we used 100 bp regions centered on the OCR summits of age-dynamic refined enhancers.

### DNA methylation in various genomic regions

Genome-wide methylation levels were calculated for each donor using the average methylation level of autosomal 10kb bins. Bins with a total cytosine count (coverage) less than 100 were excluded. For other genomic compartments (DMV, TSS+/-1k w/cgi, TSS+/-1k w/o cgi, exon (1st), intron (1st), intragenic, enhancer, and Polycomb), methylation levels were calculated as the average methylation across all autosomal regions within each compartment. DMV refers to DNA methylation valleys, TSS refers to transcription start sites, and cgi refers to CpG islands. Enhancer and Polycomb regions were defined based on our H3K27ac and H3K27me3 data, respectively.

### DNA methylation 3-state kinetic model

To estimate the time scale of the conversion between unmethylated cytosine (C), methylcytosine (mC), and hydroxymethylcytosine (hmC), we used coupled ordinary differential equations (ODEs), assuming a constant time scale across the lifespan. Since C + mC + hmC = 1, we only needed to estimate five parameters: the time constants for C, mC, and hmC (τ𝑐, τ𝑚𝑐 , τℎ𝑚𝑐), and the initial values of C and mC. These parameters were fitted to the methylation levels in age-DMR from all donors. The ODEs used are as follows:

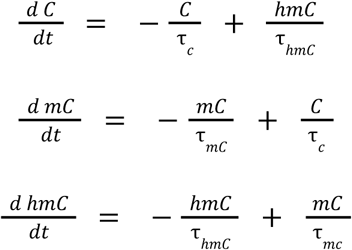

Specifically, *scipy.optimize.curve_fit* was used to fit the model to the observed methylation levels, with donor age as the independent variable. The bounds for the time constants (tau) were set between 0 and 77 (representing the maximum age of the donors), and the bounds for the initial C and mC levels were set between 0 and 1. The trf method was used for fitting, with a maximum of 8000 function evaluations.

The steady-state proportions of cytosine modifications are given by:

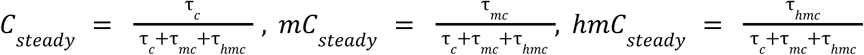

We clustered age-DMRs according to the fitted rate constants, specifically focusing on how methylation levels evolved from their initial state to their predicted steady-state, indicating whether they increased or decreased.

Furthermore, we identified the age-DMRs with τ_𝑚𝑐_ less than 5 years as “fast” groups, and those with slower rate constants as “slow” groups.

### Methylation in inactivated and escape genes

To analyze DNA methylation in X-inactivated and escape genes, we used annotations from a prior study that examined multiple human tissues^54^. We overlapped these annotations with our gene expression data, ensuring we only included genes with an average expression level greater than 4 CPM. In total, we included 295 X-inactivated genes and 51 escape genes.

---

**Supplemental Figure 1.**
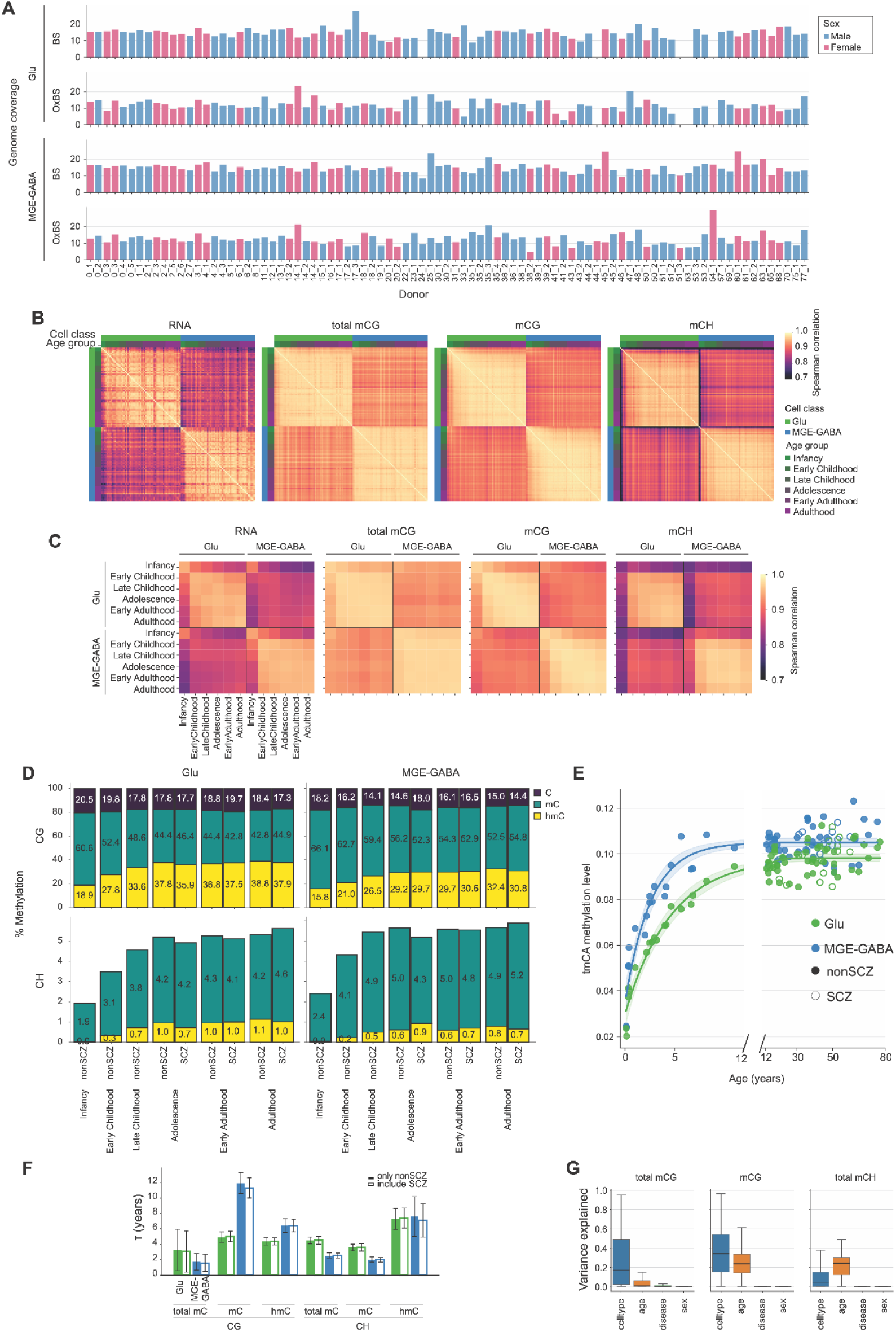
Correlation between samples and genome-wide DNA methylation in each age group; related to Figure 1. **(A)** Genome-wide coverage of individual DNA samples. **(B-C)** Spearman correlation between donors (B) and between age groups (C) in RNA expression and gene body total mCG, mCG and mCH. Expressed autosomal protein coding genes with length >1 kb were included (11,160 genes). The correlation between transcriptomes was calculated using normalized gene counts (log_2_CPM). The correlation of DNA methylation was calculated using gene body methylation level. **(D)** Mean genome-wide CG and non-CG (CH) methylation levels in each age group for Glu and MGE-GABA. **(E)** Mean genome-wide tmCA methylation of each donor for Glu and MGE-GABA **(F)** The time constant, τ, the rate of genome-wide methylation change over time, modeled by 𝑦(𝑡) = 𝑎𝑒^−𝑡/τ^ + 𝑏. Error bars represent the standard deviation of the estimate. **(G)** Variance partitioning of the mean autosomal methylation for each modality by cell type, age, disease, and sex.

**Supplemental Figure 2.**
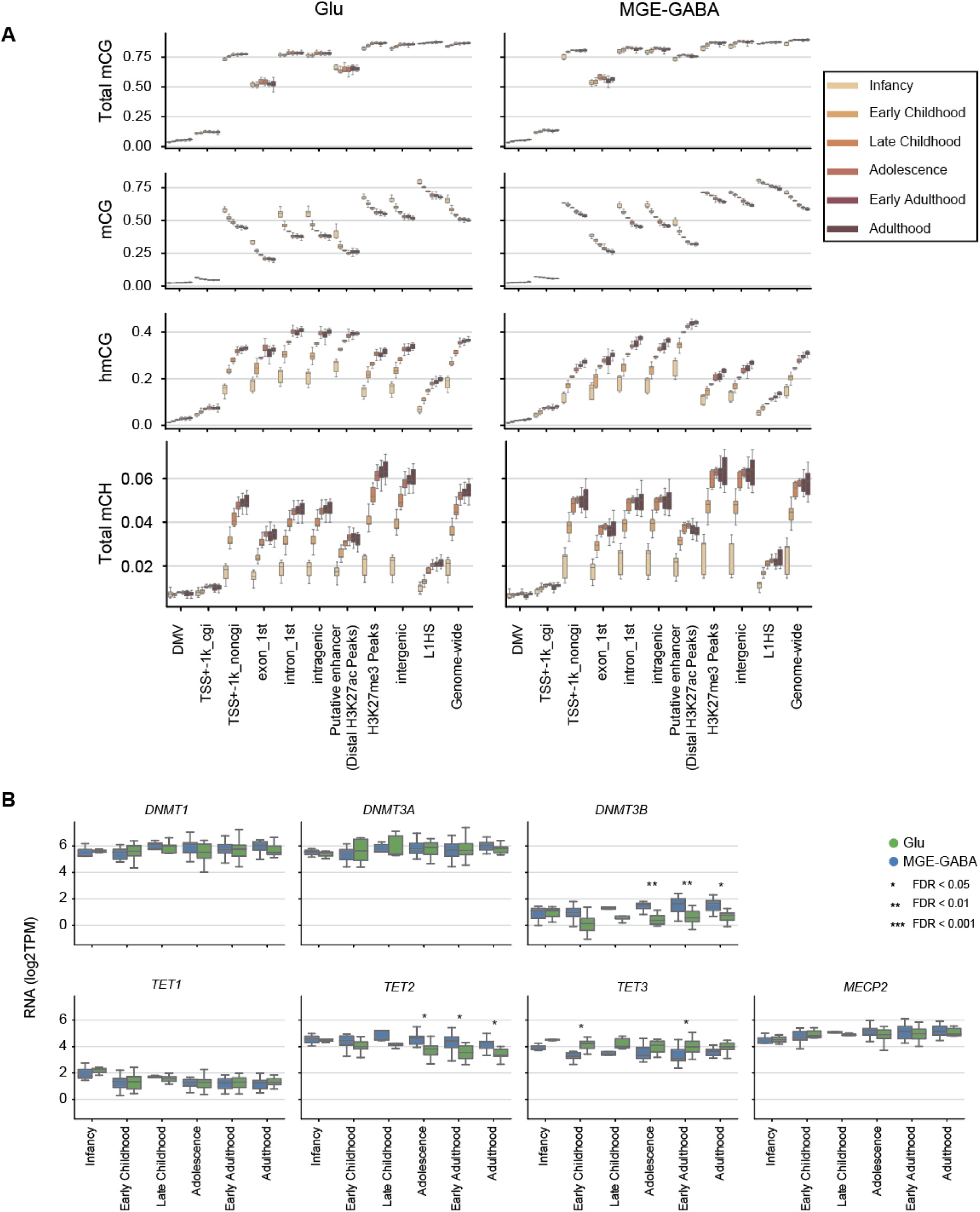
Genome-wide DNA methylation and expression of DNA methyltransferases; related to Figure 1. **(A)** Methylation levels (total mCG, mCG, hmCG) across genomic compartments and developmental stages in Glu and MGE-GABA neurons. Boxplots show methylation distribution across donors, grouped by age group. DMV: DNA methylation valleys. TSS: transcription start site. CGI: CpG island. **(B)** mRNA expression of genes encoding DNA methyltransferases and TET proteins (log2 transcripts per million (TPM)). Statistical significance between cell types was evaluated using the Wilcoxon rank-sum test.

**Supplemental Figure 3.**
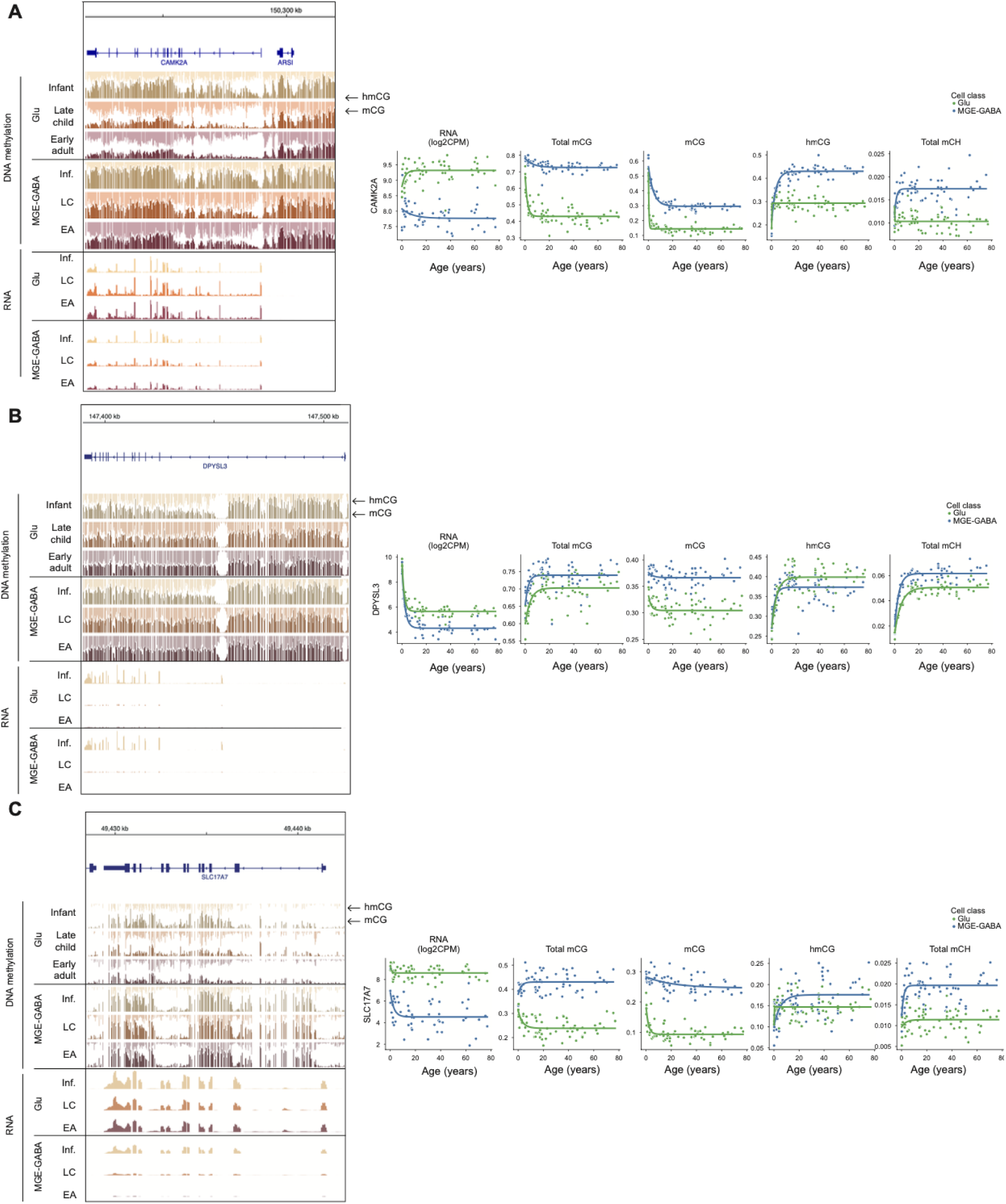
Examples of genes showing dynamic DNA methylation patterns across the lifespan related to Figure 1. Integrated genome viewer (IGV) showing the transcriptomic and epigenomic data around *CAMK2A* **(A)**, *DPYSL3* **(B)** and *SLC17A7* **(C)** in Glu and MGE-GABA neurons.

**Supplemental Figure 4.**
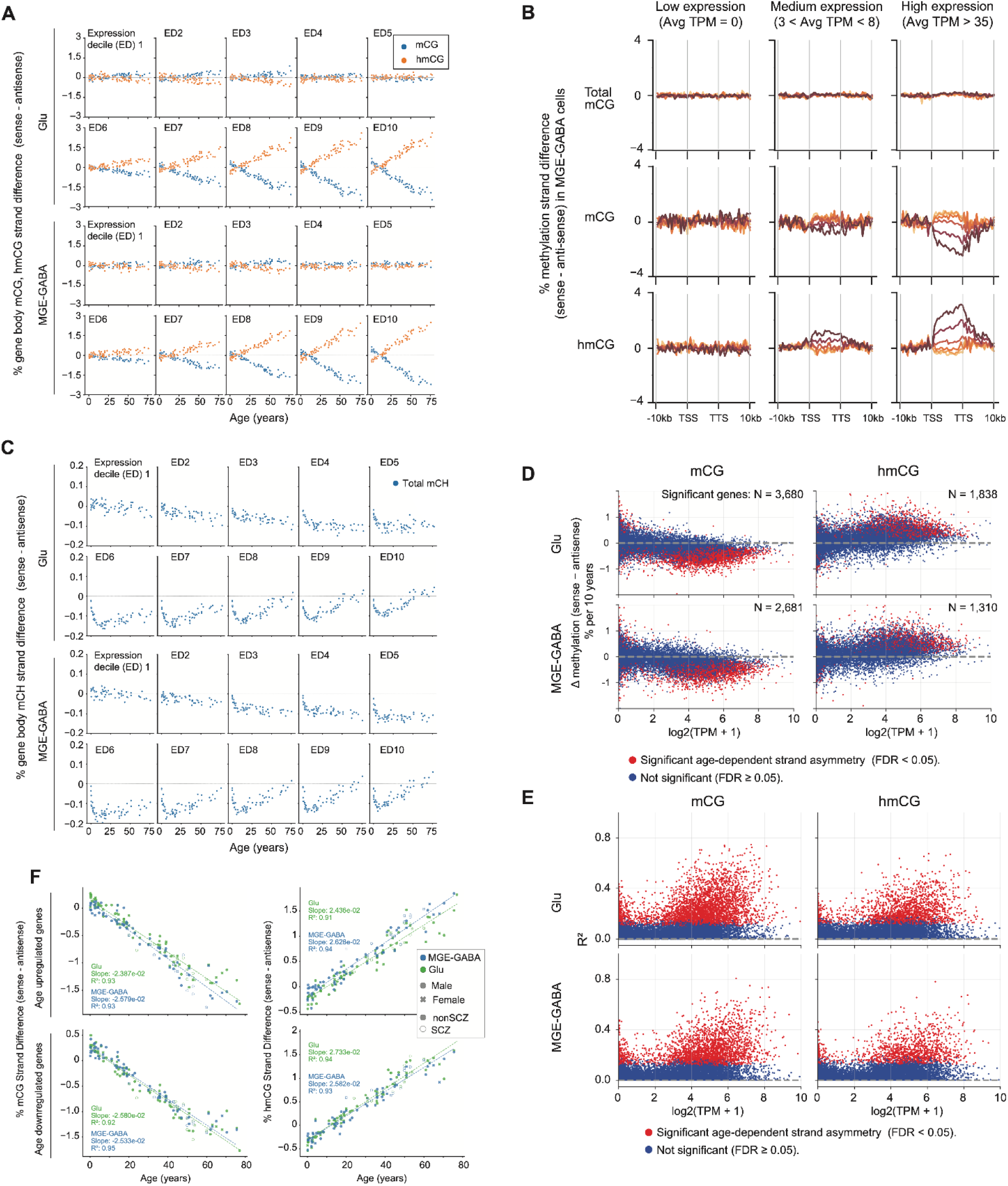
Methylation strand asymmetry; related to Figure 2. **(A)** Scatter plots of sample age versus average gene body mCG and hmCG methylation strand asymmetry, for genes grouped by into 10 expression deciles (ED1-ED10). **(B)** Metagene plots showing strand-specific differences in mCG and hmCG for low, medium, and high expression genes across age groups in MGE-GABA neurons. **(C)** Sample age versus average gene body total mCH strand asymmetry, stratified by expression decile (ED), in Glu and MGE-GABA neurons. **(D)** Scatter plots of gene expression (log₂(TPM + 1)) versus the fitted age-slope of gene-body strand asymmetry in mCG or hmCG for each gene in Glu and MGE-GABA neurons. Genes with significant age-dependent changes in strand asymmetry (FDR < 0.05) are highlighted in red. **(E)** As in (D), but showing gene expression versus model fit (R²) for each gene. **(F)** Average mCG and hmCG strand asymmetry for age-upregulated and age-downregulated genes in Glu and MGE-GABA. Linear regressions for each cell type are shown as dashed lines.

**Supplemental Figure 5.**
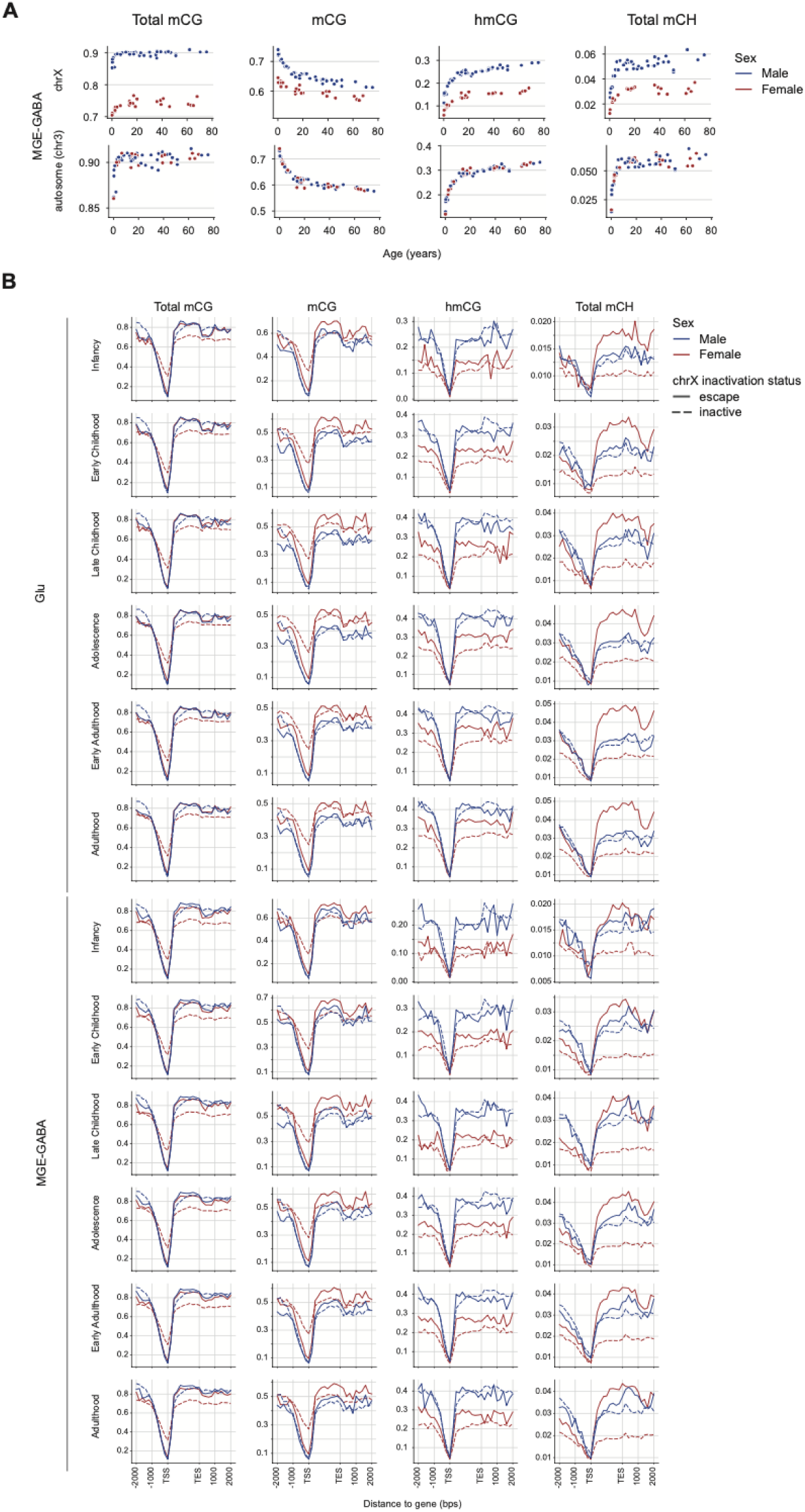
Sex differences in DNA methylation are consistent across age; related to Figure 3. **(A)** Methylation levels on autosomes and chromosome X across developmental stages in males and females in MGE-GABA. **(B)** Methylation of inactivated and escaped genes on X chromosome; shown in both sexes across age groups. TSS: transcription start site. TES: transcription end site.

**Supplemental Figure 6.**
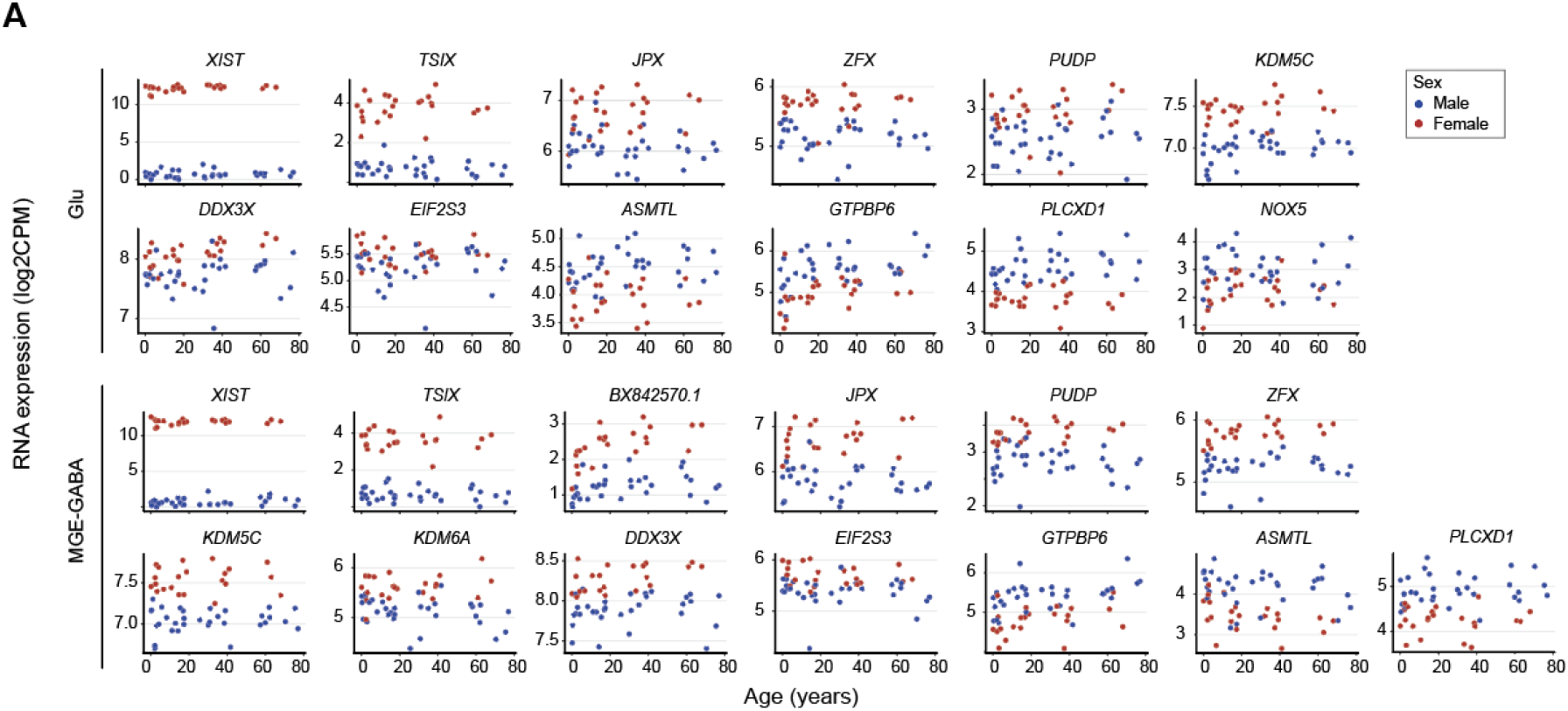
Sex-specific gene expression; related to Figure 3. **(A)** Expression of sex-differentially expressed genes (sex-DEGs) across lifespan.

**Supplemental Figure 7.**
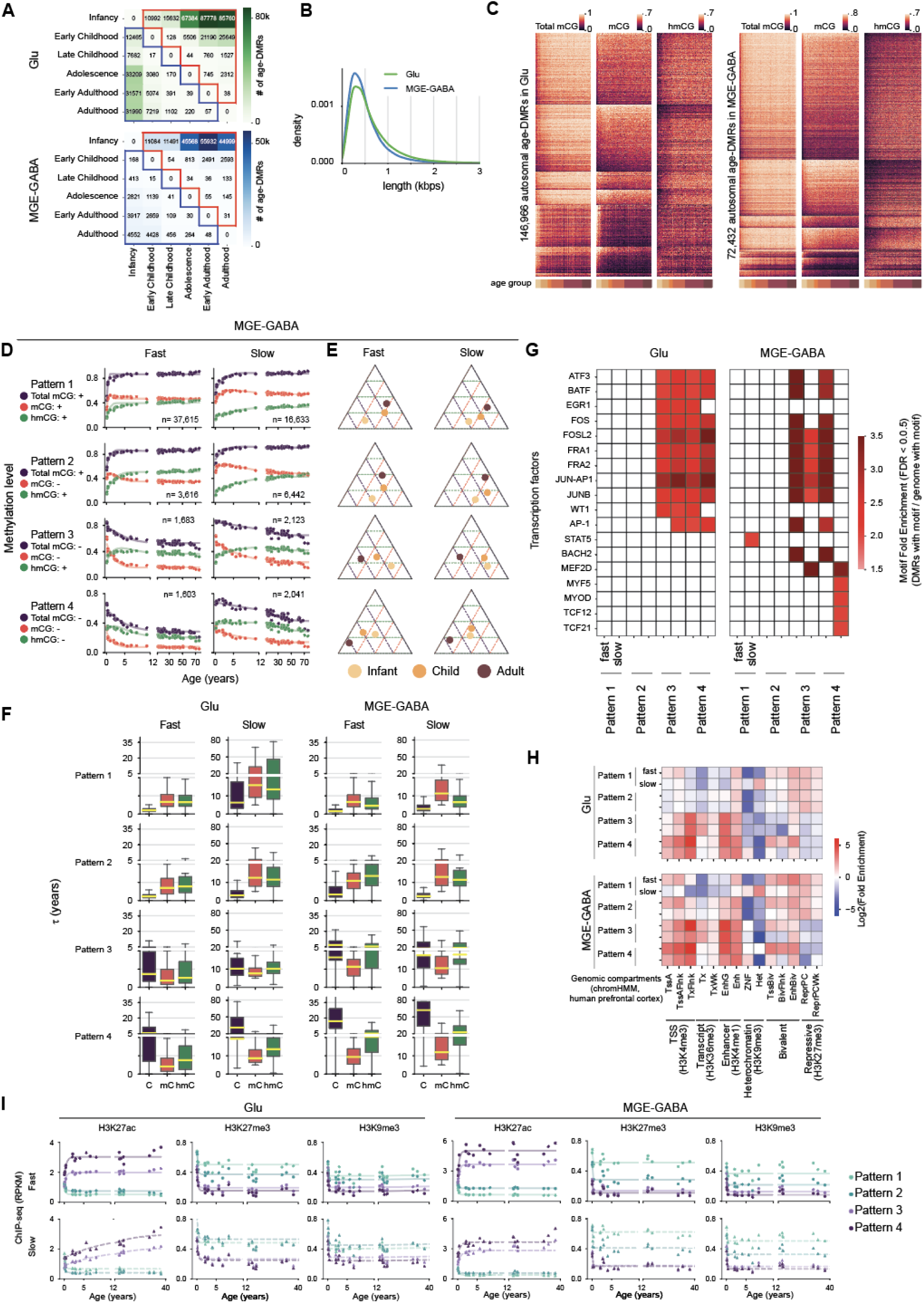
Demethylation age-DMRs are enriched in active regions with synaptic related function, related to Figure 4. **(A)** Number of age-related differentially methylated regions (age-DMRs) in pairwise age group comparisons from bisulfite sequencing (BS). The upper triangle shows the number of hypermethylated DMRs (younger < older); the lower triangle shows the number of hypomethylated DMRs (younger > older). **(B)** Distribution of the length of age-DMRs. **(C)** DNA methylation levels across all age-associated DMRs, grouped by defined age-related dynamic patterns, with donor samples ordered by age.**(D,E)** Average DNA methylation profiles of age-DMRs grouped into eight clusters based on distinct methylation dynamics (see Methods) in MGE-GABA. **(F)** Time constants (τ) estimated from the 3-state model for each cluster of age-DMRs. **(G)** Enrichment of transcription factor motifs in age-DMRs for each pattern, identified using HOMER^62^. Transcription factors with FDR < 1e-2, % of Target Sequences with Motif > 5 and enrichment >2 are shown. **(H)** Enrichment of age-DMRs in ChromHMM-defined chromatin states^103^. TssA: Active transcription start site (TSS). TssAFlnk: Flanking active TSS. TxFlnk: Transcribed state at 5’ and 3’ end of genes. Tx: Strong transcription. TxWk: Weak transcription. EnhG: Genic enhancers. Enh: Enhancers. ZNF: Zinc finger protein genes. Het: Heterochromatin. TssBiv: Bivalent TSS. BivFlnk: Flanking bivalent TSS/Enh. EnhBiv: Bivalent enhancer. ReprPC: Repressed Polycomb. ReprPCWk: Weak repressed PolyComb. **(I)** Average H3K27ac, H3K27me3 and H3K9me3 signals (RPKM) in age-DMR clusters.

**Supplemental Figure 8.**
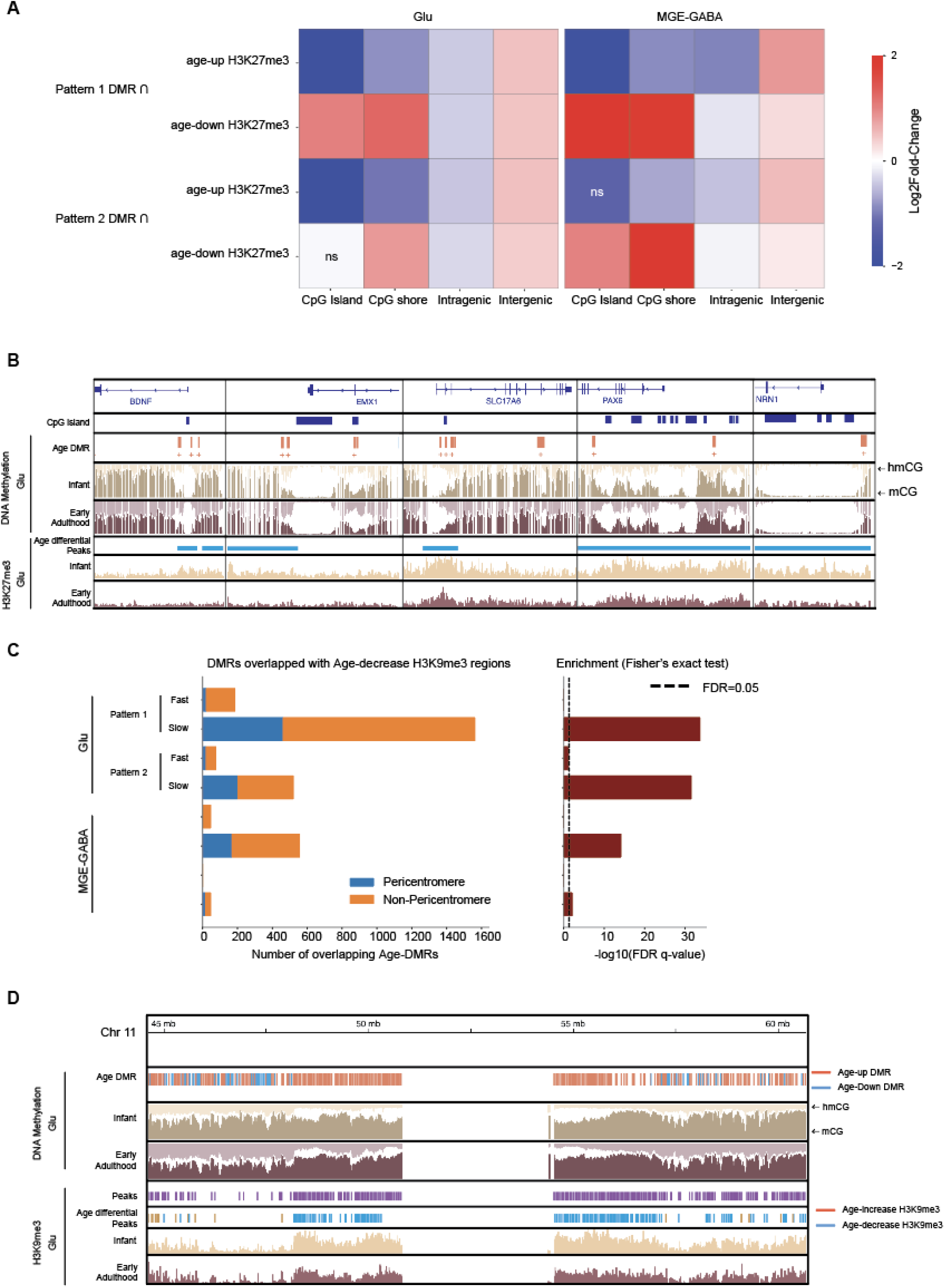
Age Differential methylated regions and Age-dependent H3K27me3 regions and H3K9me3 regions, related to Figure 4. **(A)** A genome-wide shuffling test reveals enrichment of overlapping age-DMRs and age-dependent H3K27me3 peaks within CpG islands, CpG shores, intragenic (no TSS) and intergenic (no TSS) in Glu and MGE-GABA neurons. **(B)** Progressive accumulation of hmCG from infancy to adulthood leads to high total mCG in these genes, replacing early-life Polycomb occupancy. **(C)** Number of Pattern 1 and Pattern 2 age-DMRs overlapping with age-decrease H3K9me3 regions, stratified into pericentromeric and non-pericentromeric regions. Enrichment in pericentromeres was assessed using Fisher’s exact test, with p-values adjusted for multiple comparisons using the FDR method. **(D)** Genome browser view of the pericentromeric region of chromosome 11 showing increased total mCG and decreased H3K9me3 patterns.

**Supplemental Figure 9.**
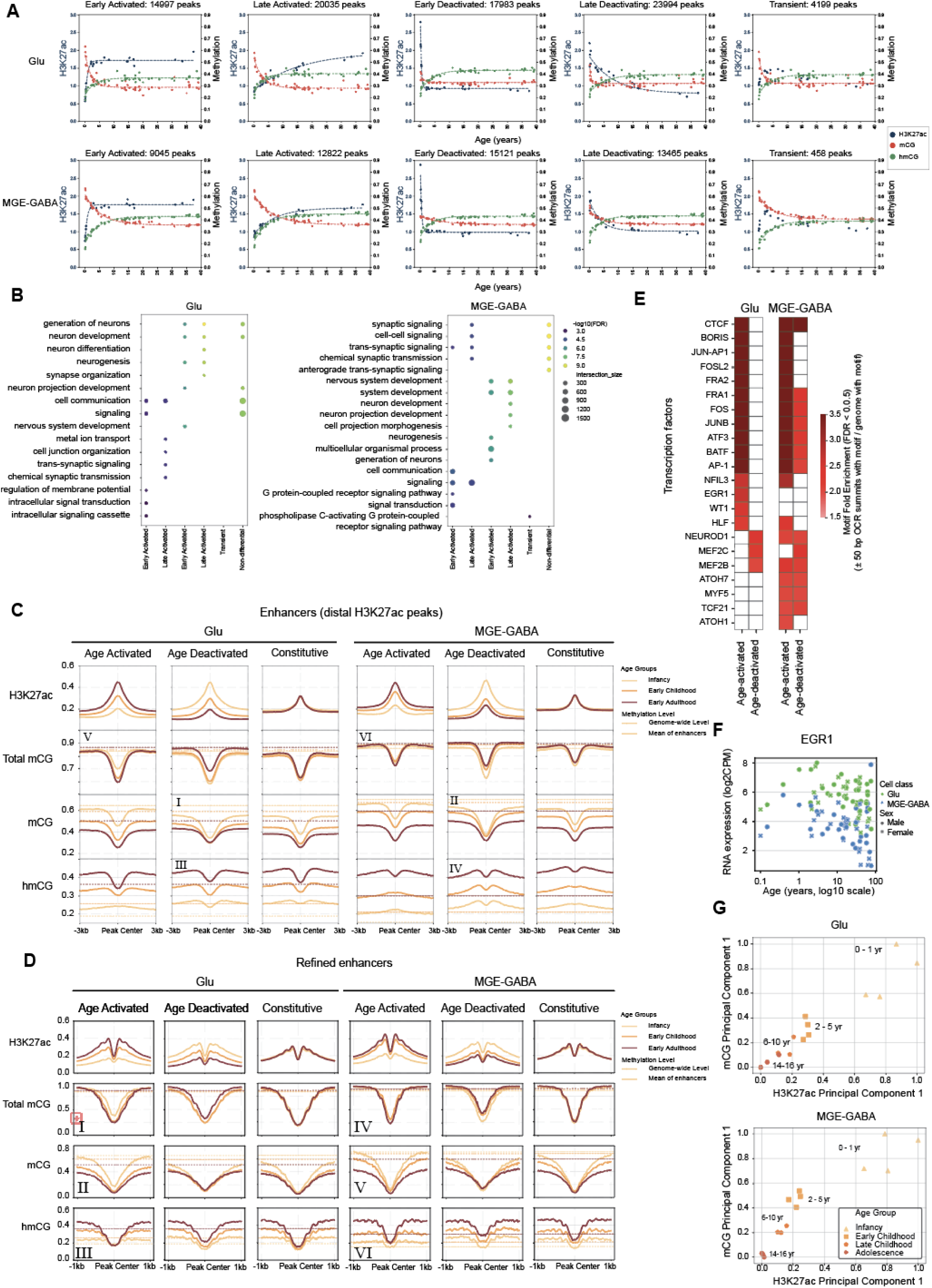
Age Differential methylated regions and Age-dependent H3K27m3 regions, related to Figure 5. **(A)** Scatter plot of individual age versus average H3K27ac (black), mCG (red), and hmCG (green) levels across five groups of age dependent H3K27ac peaks: fast activated, gradual activated, fast deactivated, gradual deactivated, and transient, shown for both Glu and MGE-GABA neurons. **(B)** Functional enrichment analysis of genes linked to distal age-activated, age-deactivated, and constitutive enhancers in Glu and MGE-GABA neurons. Enhancers were assigned to the nearest expressed gene, and enrichment was performed on the linked gene sets. **(C)** Meta-peak plot showing H3K27ac, total mCG, mCG, and hmCG signals within ±3 kb of enhancer centers, divided into activated, deactivated, and constitutive groups, for both Glu and MGE-GABA neurons. **(D)** Same as (C), but with profiles centered on refined age-dependent enhancer summits and within ±1 kb. This recentering enables assessment of DNA methylation levels within open chromatin regions (OCRs) and enrichment of transcription factor motifs in age-DMRs for each pattern, identified using HOMER **(F)** Gene expression profiles of *EGR1*.**(G)** Principal component analysis reveals coordinated developmental trajectories of H3K27ac and mCG.

**Supplemental Figure 10.**
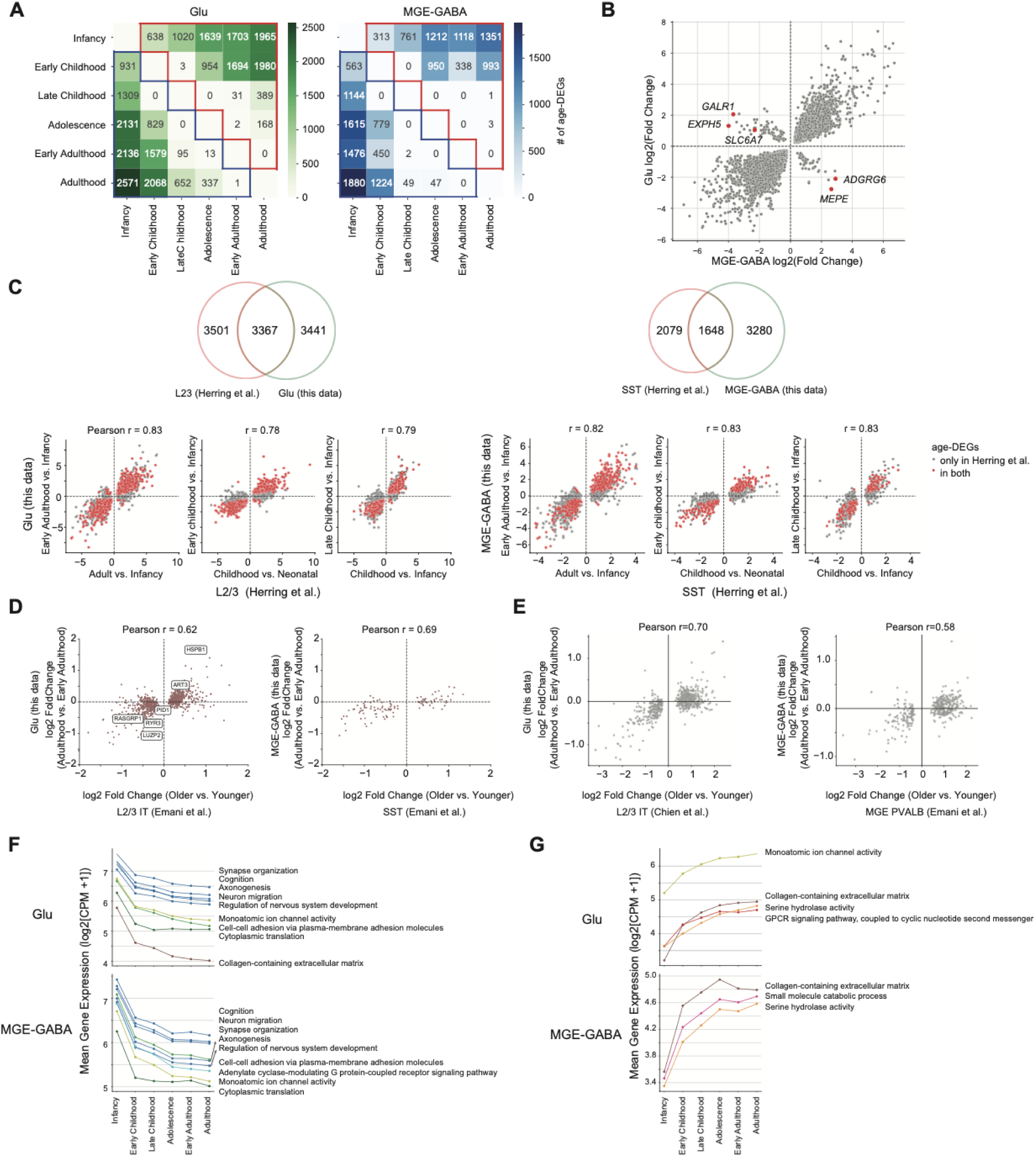
Consistency of age-DEGs with other published large scale datasets, related to Figure 6. **(A)** Number of age-related differentially expressed genes (age-DEGs, FDR < 0.05) in pairwise age group comparisons. The upper triangle shows the number of upregulated genes (younger < older); the lower triangle shows the number of downregulated genes (younger > older). **(B)** Scatter plot showing the log₂ fold-change of each gene between Glu and MGE-GABA from matched age group comparisons. **(C-E)** age-DEGs compared with published dataset from Herring et al.^78^ (C), Emani et al.^77^ (D), and Chien et al.^29^ (E). **(F)** Downregulated pathways: synaptic, neurite, adhesion, translation, ion transport. **(G)** Upregulated pathways: ECM remodeling, serine hydrolases.

**Supplemental Figure 11.**
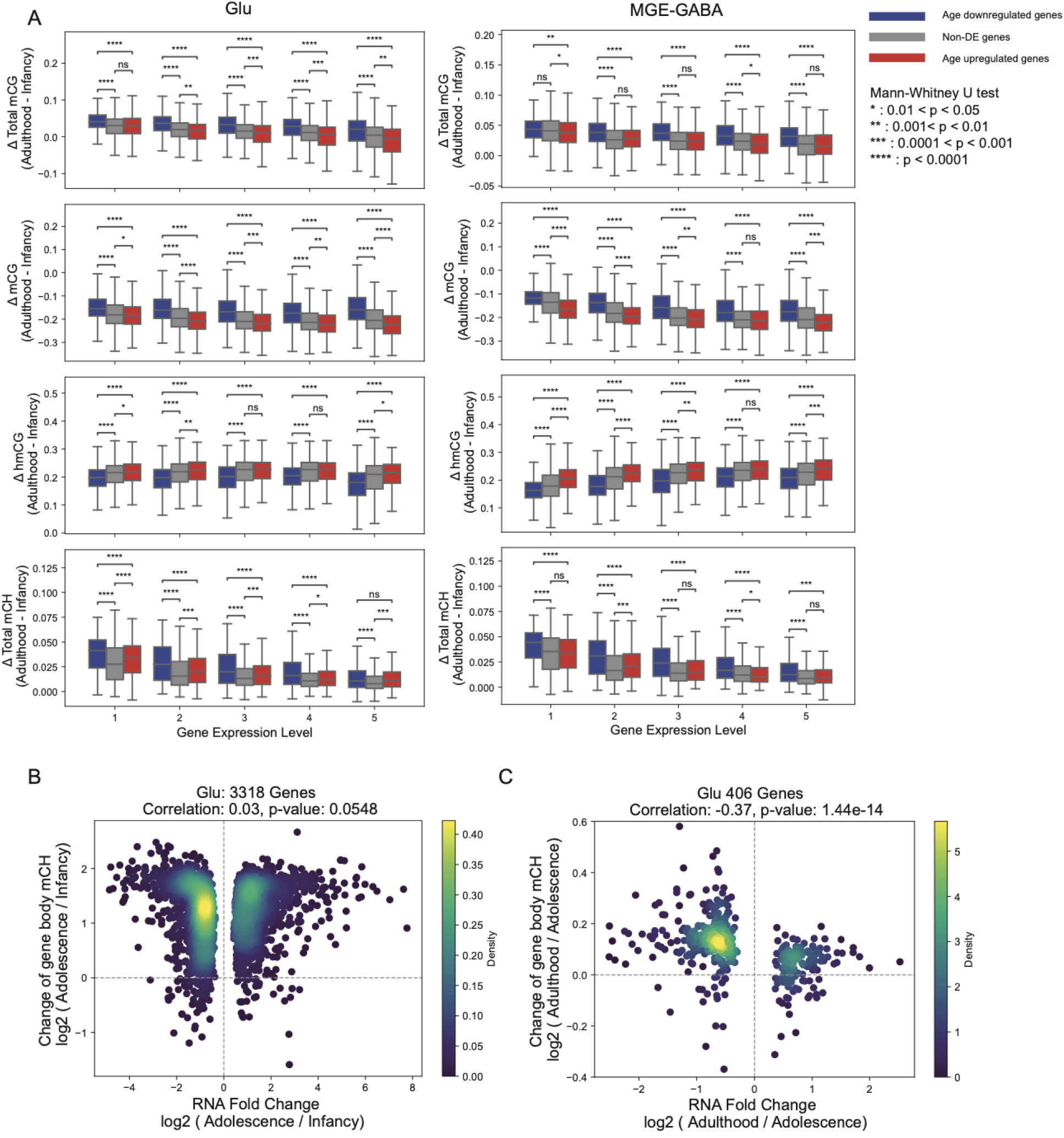
Gene body methylation change in age differential genes, related to Figure 6. **(A)** Age-related changes in total mCG, mCG, hmCG, and total mCH levels for age-upregulated, age-downregulated, and non-differentially expressed genes, stratified into quintiles based on infant expression level within each cell type. **(B)** Scatter plot showing RNA fold change between infancy and adolescence versus change in gene body total mCH for genes differentially expressed between infancy and adolescence in Glu neurons. **(C)** Same as (B), but for genes differentially expressed between adolescence and adulthood in Glu neurons.

